# Corticothalamic Projections Gate Alpha Rhythms in the Pulvinar

**DOI:** 10.1101/2021.09.10.459796

**Authors:** Nelson Cortes, Reza Abbas Farishta, Hugo Ladret, Christian Casanova

## Abstract

Two types of corticothalamic (CT) terminals reach the pulvinar nucleus of the thalamus, and their distribution varies according to the hierarchical level of the cortical area they originate from. While type 2 terminals are more abundant at lower hierarchical levels, terminals from higher cortical areas mostly exhibit type 1 axons. Such terminals also evoke different excitatory postsynaptic potential dynamic profiles, presenting facilitation for type 1 and depression for type 2. As the pulvinar is involved in the oscillatory regulation between intercortical areas, fundamental questions about the role of these different terminal types in the neuronal communication throughout the cortical hierarchy are yielded. Our theoretical results support that the co-action of the two types of terminals produces different oscillatory rhythms in pulvinar neurons. More precisely, terminal types 1 and 2 produce alpha-band oscillations at a specific range of connectivity weights. Such oscillatory activity is generated by an unstable transition of the balanced state network’s properties that it is found between the quiescent state and the stable asynchronous spike response state. While CT projections from areas 17 and 21a are arranged in the model as the empirical proportion of terminals types 1 and 2, the actions of these two cortical connections are antagonistic. As area 17 generates low-band oscillatory activity, cortical area 21a shifts pulvinar responses to stable asynchronous spiking activity and vice-versa when area 17 produces an asynchronous state. To further investigate such oscillatory effects through corticothalamo-cortical projections, the transthalamic pathway, we created a cortical feedforward network of two cortical areas, 17 and 21a, with CT connections to a pulvinar-like network. With this model, the transthalamic pathway propagates alpha waves from the pulvinar to area 21a. This oscillatory transfer ceases when reciprocal connections from area 21a reach the pulvinar, closing the cortico-thalamic loop. Taken together, results of our model suggest that the pulvnar shows a bi-stable spiking activity, oscillatory or regular asynchronous spiking, whose responses are gated by the different activation of cortico-pulvinar projections from lower to higher-order areas such as areas 17 and 21a.

## 1 Introduction

Integrating different visual attributes of an image into a single neuronal representation is a difficult task. Throughout evolution, the mammalian visual cortex has solved this computational problem by separating these different features into distinct and parallel processing modules [Bullier, 2001]. Interactions between these modules are hierarchical; from lower levels of the organization, more complex levels are created [Felleman and Van Essen, 1991, Bullier, 2003, Hegde and Felleman, 2007]. This feedforward pathway (from the retina to the lateral geniculate nucleus in the thalamus, LGN, and from here to the striate visual cortex, area 17 or V1) is accompanied by feedback projections that shape responses to those upcoming signals. Thus, visual processing consists of cortical signals traveling from lower to higher-order (HO) areas and vice-versa throughout cortico-cortical connections, whose organization follows a hierarchical pattern. Such anatomical and functional arrangement of cortical visual areas connected by specific laminar projections is the core of the cortico-centric view of visual integration [Nassi and Callaway, 2009].

Besides direct communication of cortico-cortical areas via feedforward and feedback connections, indirect communication through cortico-thalamo-cortical connections also occurs. This transthalamic pathway enables several ranges of cortical communication by single synapses in higher-order (HO) thalamic nuclei [Sherman and Guillery, 2002, Casanova 2004]. Such is the case for the pulvinar, the most prominent HO nuclei in large mammals [Shipp, 2003, Sherman and Guillery, 2011]. Interesting, neuronal responses in the pulvinar are driven mainly by reciprocal connection to almost, if not all, visual cortical areas; albeit the pulvinar receives direct connections from subcortical structures as the superior colliculus (SC) and the retina [Casanova, 2003, Casanova, 2021]. This exquisite circuitry between the pulvinar and the cortex has been recently suggested to mediate the temporality of cortical communication [Saalmann et al., 2012, Fiebelkorn et al., 2019, Cortes et al., 2020]. The pulvinar may regulate cortical responses in feedforward and feedback directions by synchronizing distant oscillatory cortical regions, given its strategic position within the visual hierarchy. Such temporal control may be crucial for shaping whole-brain dynamics on a moment-to-moment basis when, for example, attention demands or visual contrast modulation are required [Shipp, 2004, Snow et al., 2009, Cortes and van Vreeswijk, 2012, de Souza et al., 2020, Cortes et al., 2020].

Although the transthalamic pathway has been highlighted as an essential part of the visual system, how visual processing along this pathway differs from that of cortico-cortical pathways has remained elusive [Casanova, 2003]. Attempts have been made to clarify projection differences based on their pathways’ anatomical and functional characteristics. Two types of corticothalamic (CT) projections have been recognized in thalamic nuclei, including the pulvinar. Type 1 axons are thin and have long, thin branches with small terminal endings and are considered to be equivalent to round small (RS) presynaptic terminals observed at ultrastructural level [Rockland, 2018]; type 2 axons have thicker axon diameters with clustered endings considered to be equivalent to round large (RL) presynaptic terminals. These terminals originate in different layers: while type 1 projections arise from layer 6, type 2 projections from layer 5. [Ojima et al., 1996, Vidnyánszky et al., 1994, Feig and Harting, 1998, Huppe-Gourgues et al., 2006, 2019]. In addition, based on the characterization of their excitatory postsynaptic potentials (EPSPs), type 1 and 2 CT projections display frequency-dependent facilitation and depression, respectively [Li et al., 2003]. While type 1 terminals are associated with CT projections from area 17 to LGN, type 2 terminals are more abundant in CT projections from area 17 to the pulvinar [Abbas-Farishta et al., 2020]. Taken together these findings, such pathways and their associated axonal ending types 1 and 2 appear to complement synergistically each other to fine-tune visual processing in the cortex.

Although the pulvinar receives more type 2 than type 1 axon terminals from area 17, this ratio is not fixed within the visual cortex. Type 1 endings seem to be more represented in CT connections from higher hierarchical levels [Abbas-Farishta et al., 2020]. For instance, in cats, CT terminals emerging from HO cortical areas, as areas 21a (considered to be a homolog of primate area V4 [Payne, 1993]) and the posteromedial lateral cortex (PMLS, the homolog of area MT in primates ([Payne, 1993]) [Huppe-Gourgues et al., 2019], display more type 1 terminals. Furthermore, in the anterior ectosylvian visual area (AEV), one of the highest areas in the hierarchical organization of the visual system, the proportion of type 1 endings highly dominate the CT pathway toward the pulvinar. These findings suggest that the ratio of type 1/type 2 cortico-pulvinar projections seems to vary according to the hierarchical position of the source cortical area.

Such variation of CT projections along the cortical hierarchy raises questions about their functionality and, consequently, the role that pulvinar might play with cortex in visual processing. Fundamentally, as theoretical works suggest, synaptic dynamics of terminal types 1 and 2 exhibit various oscillatory patterns that dominate the response of the target network by short-term facilitation or depression, respectively [Tsodyks et al., 1998]. In addition, as experimental data shows, several low-oscillatory rhythms have been detected in the pulvinar [Saalmann et al., 2012, Fiebelkorn et al., 2019, Halgren et al., 2019]. Therefore, to investigate whether CT projection types along the visual hierarchy influence pulvinar neuronal temporal responses differently, we simulated a pulvinar-like network of excitatory and inhibitory neurons receiving both terminal types 1 and 2 from two cortical like structure simulating a low and a higher ranked cortical area. The distribution of these corticopulvinar terminals were established using projection patterns of area 17 and extra-striate area 21a of cats, whose anatomy and functional connectivity in relation to the pulvinar has been well documented [de Souza et al., 2020, Cortes et al., 2020, Abbas-Farishta et al., 2020, Abbas-Farishta et al., 2021]. Corticopulvinar projections were implemented with short-term plasticity dynamics, in which terminal types 1 and 2 had facilitation and depression of their EPSP, respectively [Li et al., 2003]. Thus, connections were established to reproduce alpha-band oscillations (7.5 Hz to 12.5 Hz) in pulvinar neurons’ populations. We found that alpha rhythms in the pulvinar are generated by each cortical area separately or by the simultaneous combination of the two areas in a specific range of connectivity weights. In the first case, when each cortical area independently evokes pulvinar alpha waves, the oscillatory low-frequency activity generated by one area was changed by irregular spiking responses (asynchronous state) when the other targets the pulvinar. This property suggests that the pulvinar has a bi-stable state of oscillatory or asynchronous responses depending on the origin of the activation coming from cortico-thalamic afferent projections along the visual cortical hierarchy.

## 2 Material and Methods

### 2.1 Network Models

Three models were used in this work to simulate the neuronal pulvinar dynamics evoked by cortico-pulvinar projections (Figure 1A). Each model is an upgrade of the previous one to provide an integrative framework in which the role of cortico-thalamic projections was investigated. The first model consists of a network in the balanced excitatory-inhibitory (*E-I*) state, in which inputs are modelled as Poisson spike trains. Such inputs have types 1 and 2 dynamics, inducing Short-Term Facilitation (STF) and Short-Term Depression (STD) responses, respectively (Figure 1B-C) [Li et al., 2013]. With this model, the proportion of input synapses and their strength were varied to investigate how this network, our “proto-pulvinar”, evoked either oscillatory or asynchronous responses. The second model is similar to the previous. However, here, the “pulvinar” network was targeted by two simulated cortical areas, each of them with a combination of type 1 and 2 synapses. As in the previous model, cells were simulated as Poisson spike trains (Figure 1A2). The proportion of type 1 and 2 projections, as well as the synaptic contact of those projections to *E-I* pulvinar neurons, were determined by available empirical data from the anatomy of cortical-pulvinar cat projections (Table 1). This model was useful to identify activation dynamic ranges of thalamic neuron populations as cortical projections reach them.

**Figure 1:**
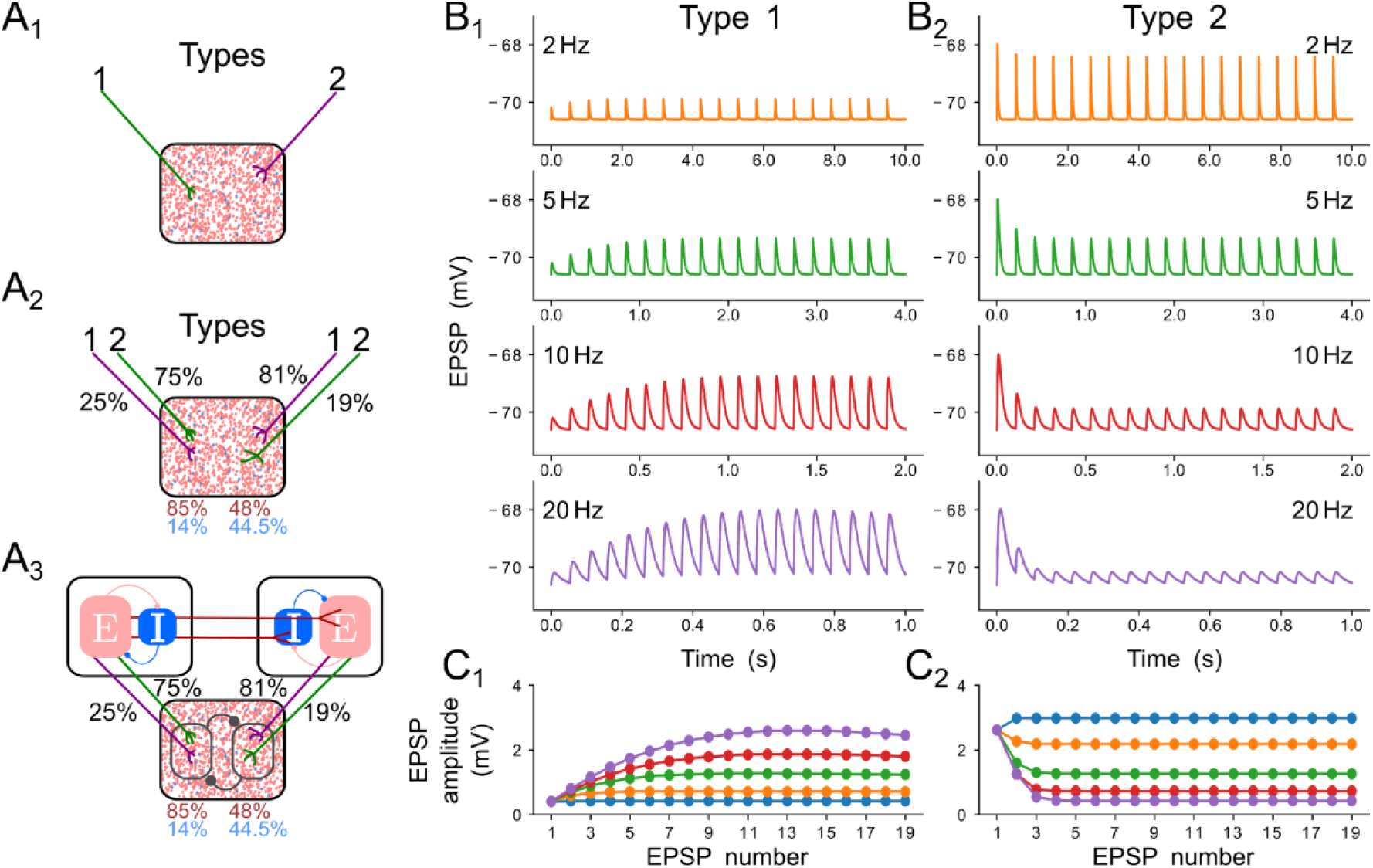
Models and short-term plasticity for synapses used in this work. A) Three models used during simulations. 1) Only terminal types 1 and 2 project to the pulvinar structure. 2) Two areas with proportion and number of contacts of terminal types 1 and 2 in areas 17 and 21a. 3) Cortico-pulvinar-cortical model consisting of three networks, a feedforward cortical pathway, a transthalamic pathway, and cortico-thalamic connections. B) EPSP of a pulvinar neuron for types 1 and 2. C) EPSP amplitude (mV) as a function of repetitions for cases 1 and 2.

**Table 1:**
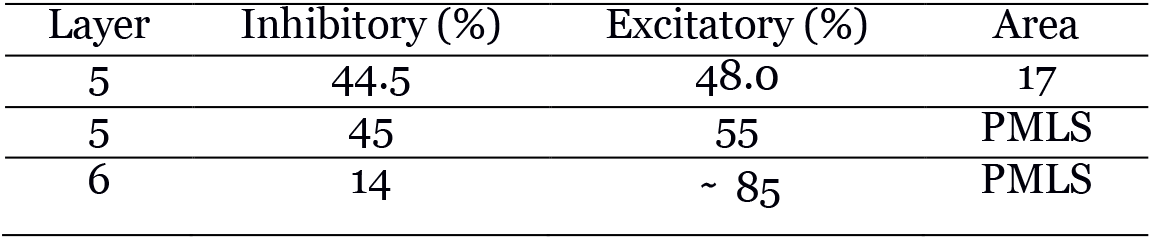
Percentage of contacts from layers 5 and 6 of areas 17 and PMLS to excitatory and inhibitory cells in the pulvinar.

| Layer | Inhibitory (%) | Excitatory (%) | Area |
| --- | --- | --- | --- |
| 5 | 44.5 | 48.0 | 17 |
| 5 | 45 | 55 | PMLS |
| 6 | 14 | ~ 85 | PMLS |

Finally, the pulvinar network was targeted by other two independent networks that imitate cortical areas 17 and 21a (Figure 1C3). Each cortical area was organized to reach the *E-I* balanced state. A feedforward connection, from areas17 to 21a, was established, and reciprocal connection between the cortices and the pulvinar-like structure were settled. Here, cortical axons terminal to pulvinar neurons were simulated with short-term plasticity dynamics, but synapses from the LGN to the area 17 (modeled as Poisson spike trains) and between areas 17 and 21a had linear integration of their synaptic inputs. Cortico-pulvinar projections with type 1 and 2 terminals were chosen randomly from neurons in areas 17 and 21a, and their proportion and contact to *E-I* pulvinar neurons were organized as in the previous model. This model does not consider direct feedforward connections from LGN to area 21a [Wimborne and Henry, 1992]. Each area, including the pulvinar network, was simulated with *N* = 10000 neurons [Vogels and Abbott, 2009, Cortes and van Vreeswijk, 2015, de Souza et al., 2020]. From this number of neurons, 20% were inhibitory cells [Rodney et al., 2004]. The effect of separated cortico-recipient zones in the pulvinar and their consequences in the response of 21a neurons were studied with this model.

### 2.2 Neuron and Synaptic Dynamics

Thalamic and cortical neurons were modeled with adaptive exponential integrate-and-fire dynamics [Brette and Gerstner, 2005]. This model consists of two coupled differential equations describing the leak current (linear component) and the spike generation component (exponential function). Also, an adaptive current, *w*, was added to simulate action potential adaptation. The membrane potential of a neuron (*i, A, β*) is given by:

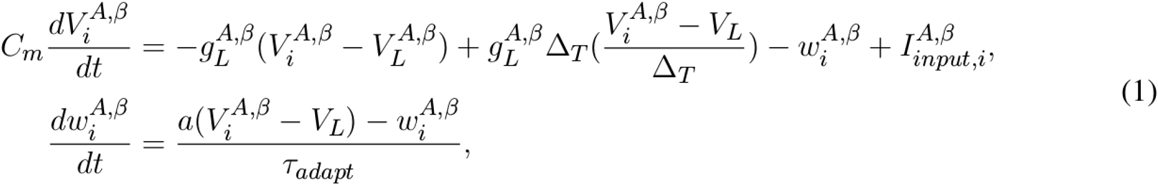

where *Cm* is the capacitance of the neuron, *V_L_* is the leak reversal potential, *V_T_* is the threshold and Δ_*T*_ is the slope factor, *τ_adapt_* is the time constant and *a* describes the level of subthreshold adaptation. Every time that the neuron *i* fires, *w* is increased by a current *b* (spike-triggered adaptation), and the membrane potential is reset to a fixed voltage, *V_r_* of the neuron, *i*, which has, *A*, excitatory or inhibitory actions, and determinated as cortical of thalamic component, *β*. Only excitatory neurons have adaptation current dynamics.

The input current that a neuron (*i, A, β*) receives is:

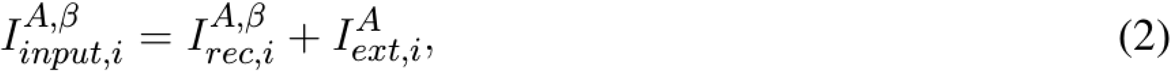

where *I^A^_rec_* characterizes the synaptic current from recurrent connections of each area. When cortical cells are integrated in a network, the external current, *I^A^* comprises one or two terms, if the unit comes from area 17, or from area 21a or pulvinar, respectively.

Synapses for cortical and pulvinar recurrent connections and from area 17 to 21a (feedforward component) are simulated as an instantaneous rise of synaptic current followed by an exponential decay.

Short-term synaptic plasticity (STP) is implemented with a phenomenological model [Stimberg et al., 2019b]. This model considers synaptic release as the product of two variables, *x_s_* and *u_s_*, where *x_s_* represents the fraction of the total neurotransmitter that remains available for release, and *u_s_* reflects the fraction of available resources ready for use, that is, the resources of neurotransmitter “docked” for release by exocytosis by calcium sensors. After an action potential and the beginning of another, *u_s_* decays to 0 at rate *ω_f_*, and *x_s_* recovers to 1 at rate *ω_d_*, as:

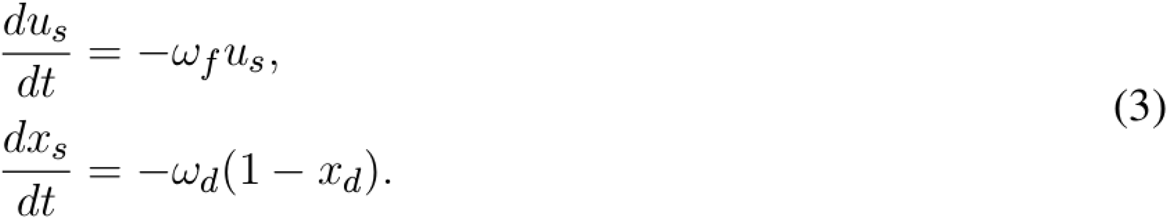

The influx of calcium in the terminal triggered by the arriving of action potentials modifies a fraction *U*_0_ of neurotransmitter resources not expected for release (1 – *u_s_*) to the “docked” state ready to be released (*u_S_*). Eventually, a release *r_s_* from the fraction of *u_s_* of the available neurotransmitter resources are generated, while *x_s_* decreased by the same quantity, so:

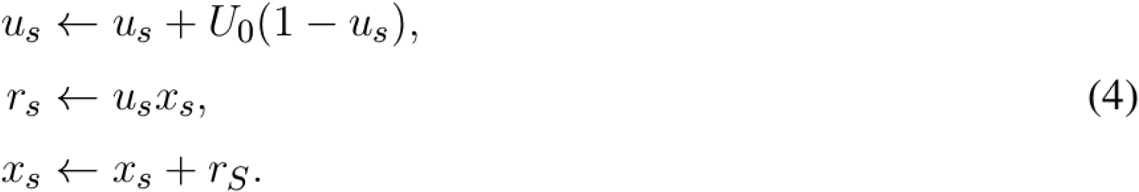

When a presynaptic action potential arrives, *A* synapses increase the *A* conductance, g^A^_STP_ in the postsynaptic neuron as g^A_STP_^ ← g^A_STP_^ + w_A_r_s_, where A = E, I. Only excitatory connections as short-term plasticity dynamics, given that only excitatory long-range cortico-cortical and cortico-pulvinar terminal have been described.

### 2.3 Feedforward and Recurrent Connections

Recurrent and external connectivity for each structure were random with connection probability, *p*, specific to *E* and *I* populations (*p_AI_=0.1, p_AE_=0.5*). Input current is defined as 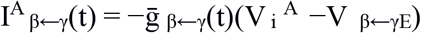, where the term on the right side of the equation is the sum of all conductance from all presynaptic inputs on the neuron (*i, A, γ*), where *γ* is the source and β the targeting structure. In general, it is described as:

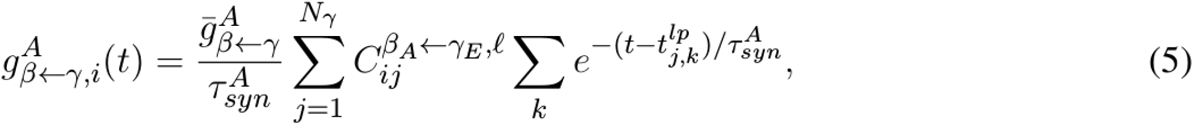

where *t^γ^_j,k_* is the time of the *k*th action potential of the neuron (*j, γ*), where *γ* can be the LGN, area 17 or the pulvinar, and *β* can be the pulvinar, area 17 or area 21a. For the recurrent connectivity in the pulvinar and the cortex, *β* = *γ*, and *i* ≠ *j*. The connection matrices *C^β_A_←γ_E_^*, for *A* = *E, I*, are random with probability *c*_*β←γ*_*K/N_β←γ_* and *C^β_A_←γ_E_^* = 0 otherwise. On average, neurons type *β* receive *K_β←γ_* = *c_β←γ_K* presynaptic connections from *γ* neurons. Here, the conductance 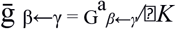 where *G^a_β←γ_^* is independent of *K*. For background synaptic activity, a population of simulated neurons (*N_bckgrnd_=8000*) as a Poisson-type spike train is applied to the pulvinar following the same synaptic dynamics of equation 5. The spike train is excitatory and activates excitatory and inhibitory neurons with a discharge rate of 0.1 sp/sec.

When invoking STP for the cortico-thalamic pathway (*γ=areas 17 or 21a, β*=pulvinar), since conductance changes over time, the voltage integration assumes an effective synaptic weight, *g^A,STP _β←γ_^* rather than the static synaptic weights *g^−A_β←γ_^*.

### 2.4 Parameters

The parameters for the cell dynamics were Cm = 1 μ F/cm2, with conductance of leak currents of gL, E = 0.1 mS/cm2 and gL, I = 0.05 mS/cm2 for excitatory and inhibitory neurons, respectively. The other parameters that characterized the dynamic of neurons with a regular spiking are: VL = −70.6 mV, V_T_ = −50.4 mV and ΔT = 2 mV. The parameters for the adaptation current were a = 24 nS, b = 0.01 nA, and τ_adapt_ = 60 ms. For bursting VL = V_T_ + 5 mV, and τ_adapt_ = 20 ms, a = 4 nS, and b = 0.5 nA. For each area, the synapses’ parameters were G_E0_ = 1.425 ms nS/cm2. G_I0_ = 1.89 ms nS/cm2, G_EI_ = 9.0 ms nS/cm2, G_II_ = 13.5 ms nS/cm2, G_EE_ = 22.5 ms pS/cm2, G_IE_ = 67.5 ms pS/cm2, with τsyn = 3 ms and V_E_ = 0 mV and V_I_ = −80 mV. For STP, parameters were separated for terminal types 1 and 2, so synaptic release probability at rest *U*_0_^type1^ = 0.006 and *U*_0_^type2^ = 0.8; synaptic depression rates *ω^type1^f* = 0.48 sec^-1^ and *ω^type2^f* = 2.0 sec^-1^; synaptic facilitation rate *ω^type1^_d_* = 1.5 sec^-1^ and *ω^type2_d_^* = 3.33 sec^-1^; and, the synaptic conductance w^A^ = G_A0_, for A = E, I. Recurrent connectivity for each area (pulvinar, areas 17 and 21a when are modelled) is K, and the probability of connection was p_A_ = K_A_/N_A_, for A = E, I.

#### 2.4.1 Variation of Pathway Connections

We used the factors *W_FF_* = 5 and *W_CP_* = 1.5 to change the weights of feedforward and cortico-pulvinar projections. These factors multiply the ratio *G*_*E*0_/*G*_*I*0_ for those entry inputs.

Simulations of network architecture and neuron equations were performed with Python version 3.2 using Brian2 simulator ([Stimberg et al., 2019a]. Euler’s integration was implemented using a time step of 0.05 ms. The accuracy of the results was verified by repeating simulations with smaller time steps (0.025 ms)

## 3 Results

As stated above, three models were used to investigate the oscillatory gating generated by cortico-thalamic terminals in the pulvinar. The first model analyzed the effect of the combination between type 1 and 2 terminals that target excitatory and inhibitory (*E-I*) neurons, whose responses are in the balanced state (Figure 1A1). This model provided a basic approximation of the weight ranges of cortico-pulvinar connections that produced oscillatory and asynchronous neuronal responses in this network, our simulated pulvinar. The second model also consisted of external projections to a pulvinar-like structure. However, this model considered two external areas, cortical areas 17 and 21a, whose cortico-pulvinar projections contain a combination of types 1 and 2 synapses (Figure 1A2). The fraction of type 1 and 2 terminals and the percentage of synaptic contact reaching *E-I* pulvinar neurons were arranged with available empirical data (Table 1). This model allowed investigating the dynamics of pulvinar responses when the activation between cortical projections from areas 17 and 21a was temporarily deferred. While the two first networks used cortical inputs as Poisson spike trains, the third model simulated explicitly cortical neurons. Here, two similar networks of *E-I* neurons were implemented and connected feedforwardly to reproduce the interaction between areas 17 and 21a (Figure 1A3). Each cortical area target pulvinar neurons, with the proportion and axon terminal contacts settled in the second model. Furthermore, for this model, cortico-pulvinar projections were divided in striate- and extrastriate-recipient zones to investigate the effect of different signalling pattering of the transthalamic pathway across cortical areas (Figure 1A3).

### 3.1 Pulvinar EPSPs Evoked by Single Cortico-Thalamic Activation

Before analyzing the complete pulvinar network, synaptic plasticity of excitatory postsynaptic potentials (EPSPs) was simulated to obtain similar qualitatively experimental magnitudes of those found in neurons of the lateral posterior nucleus [Li et al., 2003], the homologous nucleus to the pulvinar in rats. To that end, a single cortico-thalamic fiber contacting a single neuron with exponential integrate-and-fire dynamics was simulated (Mat & Met, Equation 1). Cortico-thalamic fibers were implemented with synaptic plasticity that simulates short-term facilitation (STF, Figure 1A1) and short-term depression (STD, Figure 1B1). Then, EPSPs from the pulvinar neurons were recorded as the cortical afferent fiber was stimulated with four sets of frequency impulses (0.5, 2, 5, 10, 20 Hz) (Figure 1B). With this set up, the strength of the cortico-thalamic fiber was investigated, and experimental amplitude of pulvinar EPSPs were recovered.

Two types of cortico-thalamic fiber responses were generated (Figure 1B). These simulations revealed that changes of EPSP amplitudes elicited by the stimulus train at various frequencies follows experimental results of thalamic responses at a given set of synaptic parameters. For simulated type 1 cortico-thalamic fibers, the amplitude of EPSPs enhanced as the stimulation increased in frequency, depicting a saturation in the response after seven consecutive impulses (Figure 1B1). A contrary response was seen in type 2 cortico-thalamic fibers. Here, EPSP amplitudes decreases at higher frequencies, showing a constant response of amplitude after five consecutive impulses. The amplitude of pulvinar EPSPs showed a similar profile as frequency stimulation were higher than 5 Hz. Taken together these results, our simulations evoked type 1 EPSPs with frequency-dependent facilitation (Fig 1B1), and frequency-dependent depression for type 2 EPSPs in pulvinar neurons.

### 3.2 Pulvinar Network Responses Evoked by Type 1 and 2 Terminals

The next step was to study the effect of cortico-thalamic terminals in a population of *E-I* in the balanced state (Figure 1A1). The pulvinar was modelled with sparsely connected neurons but strong connections between *E-I* populations [van Vreeswijk and Sompolinsky, 1996]. In this network, excitatory frequency-dependent type 1 and 2 terminals identified in the previous section were feedforwardly connected to excitatory and inhibitory pulvinar neurons. The weights of these external “cortical” synapses, *G*_*E*0_ and *G*_*I*0_, were invariant in time. The different responses of the pulvinar network were then analyzed when a factor *η* amplified the excitatory cortical pathway to *E-I* neurons. The logic of the *η* factor was to conserve the feedforward ratio (*G*_E0_/*G*_*I*0_) and only modulate the amplification of the external pathway to pulvinar neurons [Cortes and van Vreeswijk, 2015]. *η*_1_ and *η*_2_ were defined to amplify type 1 and 2 synaptic terminals, respectively.

#### Asynchronous and synchronous responses

Two clear states the pulvinar network showed when *η* was changed: a strong irregular (asynchronous state) or a regular (synchronous state) pattern of spiking activity (Figure 2). These two activation states were evoked with a Poisson input spike train of 10 sp/sec, in a network that had equal synaptic strengths to *E-I* populations of neurons. To characterize further both regimes, peristimulus-time histograms (PSTHs) and the statistics of spike discharges were analyzed. As Figure 2A shows, PSTHs for excitatory and inhibitory neurons reflect the global asynchronous (Figure 2A1-B1) and synchronous (Figure 2A2-B2) states of the pulvinar network. For the asynchronous state, spike statistics showed an exponential average firing rate and a normal distribution of coefficient of variation (CV) for the inter-spike interval (ISI), signatures of an irregular activation regime (Figure 2C1-D1). This was not the case for the synchronous state (Figure 2C2-D2), in which a predominant oscillatory regime in the PSTH was revealed (Figure 2B2). Here, both the average firing rate and the ISI CV had a narrow distribution profile. These differences between the two pulvinar response states were also observed when the PSTH spectrum of frequencies was measured. In the asynchronous state, a peak of oscillatory activity was observed at low frequencies (< 5 Hz), whereas, in the synchronous regime, the oscillatory activity showed a clear peak at 12 Hz, which was within the range of alpha-band oscillatory frequency. The amplitude for this peak was higher for the synchronous than the asynchronous state. In summary, an asynchronous state was characterized by a high CV (close to 1) with a low amplitude frequency peak of the PSTH, while the synchronous regime had low average firing rate and CV values, and a high oscillatory frequency amplitude of its PSTH.

**Figure 2:**
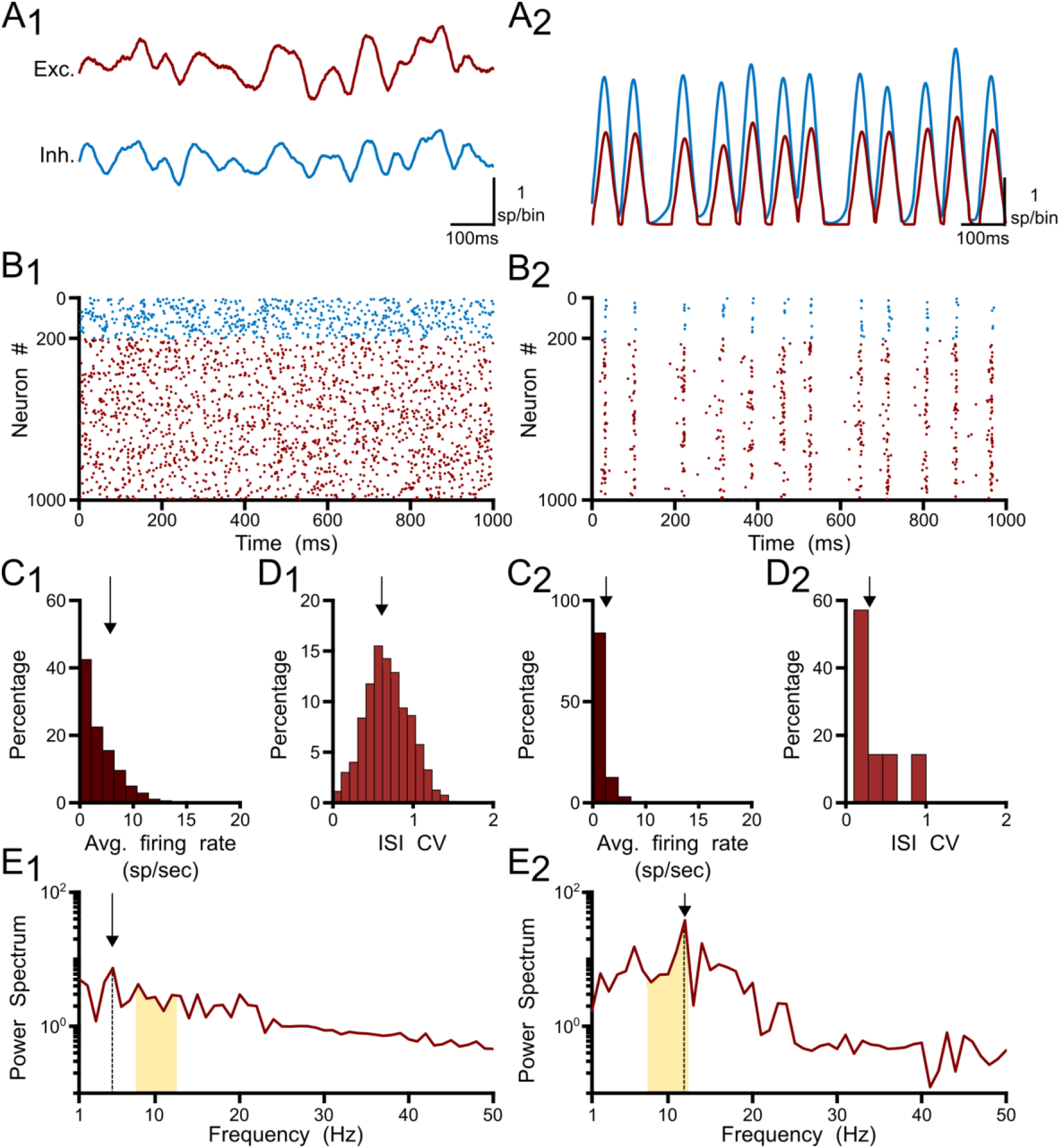
Two responses of the pulvinar network: 1) asynchronous and 2) synchronous states. A) PSTH (sp/bin) of excitatory (red line) and inhibitory (blue line) when a spike train with Poisson statistics of 10 sp/sec is applied to the network, B) Raster plot for excitatory (red) and inhibitory (blue) neurons. Only 1000 cells are shown. C) Distribution of average firing rate (sp/sec), D) Interspike interval (ISI) of the coefficient of variation (CV), E) Power spectrum of PSTHs. Arrows show the maximum frequency amplitude. Yellow zones characterize alpha-band oscillation range.

To identify in which set of values such neural states occurred, inputs and synaptic feedforward strengths to the pulvinar network were gradually increased. The variations were tested in two networks that had similar strengths of *E-I* connectivity. Figure 3 shows the result of such simulations. For type 1 terminals, the gradual increase of the input produced a gradual increase of the discharge of the neurons, with a clear asynchronous state of the network (Figure 3A1-B1). This gradual increase also occurred when *η*_1_ increased, in which the average firing rate and CV increased further (Figure 3C1-D1). Another scenario was observed for type 2 connections. Here, the pulvinar network showed a synchronous transition between the inactive and asynchronous states (*η*_1_ ~3). During this transition, the firing rate as well as the oscillation amplitude increased to high value inputs, while their CV was low. Note that this oscillatory transition also occurred for type 1 synapses, but this was less pronounced. In fact, type 2 terminals evoked the oscillatory transition at lower connection intensity strengths than those simulated for type 1 connections.

**Figure 3:**
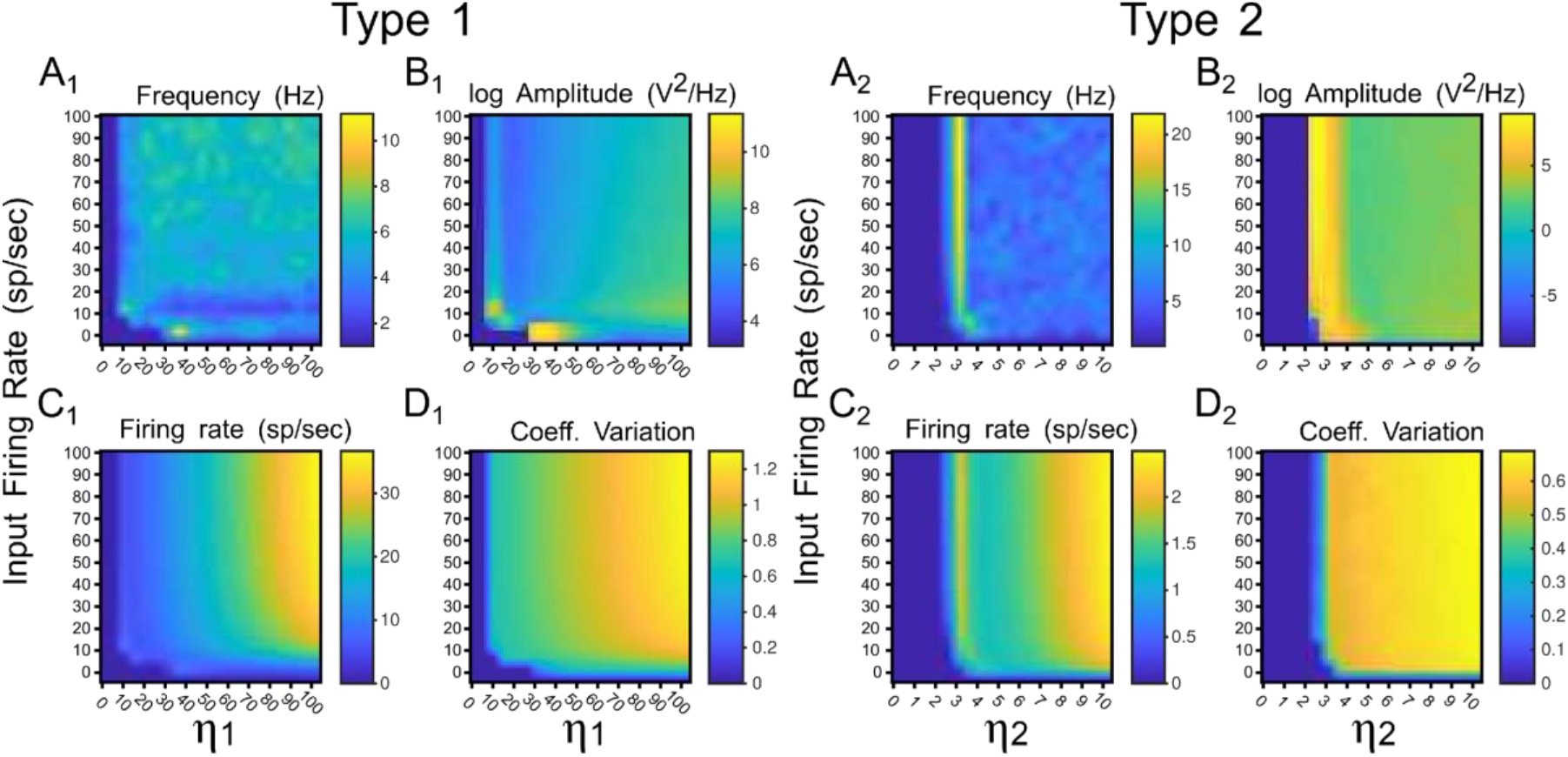
Quantitative outputs for a pulvinar network whose neurons receive terminal types 1 (1) and 2 (2) as input, and connection weights (*η*i and *η*2, respectively) increase gradually. A) Frequency (Hz), B) Amplitude (v^2^/Hz), C) Firing rate (sp/sec), D) CV.

### 3.3 Oscillatory Responses Evoked in the Pulvinar by Lower and Higher Cortical Areas

While the distribution of cortico-pulvinar terminal types 1 and 2 seems to be hierarchical-level-dependent, the proportion of contacts to excitatory and inhibitory neurons seem to be terminal-type-dependent. For area 17, the distribution of terminal types 1 and 2 are 25% to 75% respectively. In higher cortical levels, such as area 21a, this distribution is almost reversed in which types 1 and 2 are 81% and 19% of the cortico-pulvinar connections, respectively [Abbas-Farishta et al., 2020]. On the other hand, cortico-pulvinar type 1 terminals contact 85% and 14% of excitatory and inhibitory cells, respectively, type 2 synapses 48% and 44.5%, respectively [Vidnyánszky et al., 1994]). In fact, round large (RL) terminals from area 17 to the pulvinar are predominantly located in the striaterecipient zone. For extrastriate cortical areas, terminals zones are characterized as small boutons (RS) [Huppe-Gourgues et al., 2006]. Thus, cortico-thalamic projections seem to exert different synaptic actions depending on their origin along the cortical visual hierarchy and the type of terminals contacting pulvinar neurons.

Such anatomical attributes were implemented in the following simulations of the pulvinar net-work. For area 17, contributions from a Poisson spike train were divided in terminal types 1 and 2, with the number of implicit excitatory cells considered in the feedforward cortico-thalamic pathway being 25% and 75% of *K*, respectively, where *K* is the average number of total projections. For area 21a, the percentage for types 1 and 2 terminals were 81% and 29% of *K*, respectively. Regardless of their cortical area of origin, type 1 terminals had a connection probability with excitatory and inhibitory pulvinar neurons of *p* = 0.85 and *p* = 0.14, respectively, and type 2 terminals probability contacts were *p* = 0.48 and *p* = 0.445, respectively. To that network, *η*_1_ and *η*_2_ were increased gradually and input of 10 sp/sec was applied to analyze the performance of the pulvinar network. Representative frequency, amplitude of this frequency, firing rate and CV were collected after 1 sec of simulation, and are depicted in Figure 2.

#### Effect of single cortical activation

Regardless of whether the input came from area 17 or 21a, at a given strength of the feedforward cortical pathway, the pulvinar network evoked oscillations in the frequency range of 7.5-12.5 Hz (alpha waves). These oscillations were generated at the transition between the quiescent and the asynchronous steady state of the network. In the transition, the network showed a maximum frequency of ~25 Hz (beta-waves). Alpha frequency bands were found besides such maximum frequency oscillation (Figure 4, red lines). Alpha waves were symmetrical at the border of the beta bands, but for *η*_2_ high and *η*_1_ low, only one side of the transition showed alpha band responses. For areas 17 and 21a, alpha waves were around *η*_1_ = 13 and *η*_2_ = 3, and *η*_1_ = 7 and *η*_2_ = 5, respectively. Between these strengths, decreasing and increasing magnitudes of *η*_1_ and *η*_2_, respectively, allowed a continuous transition where alpha waves were always presented. Note that in order to reach such transition threshold, *η*_1_ was higher in area 17 than in area 21a. The inverse happened for *η*_2_, whose magnitude was lower in area 17 than in area 21a. Such transition threshold indicates that the proportion and the distribution of synaptic contacts of terminal types 1 and 2 settled for areas 17 and 21a promote lower values of the cortico-pulvinar connection to reach oscillatory alpha-band activity.

**Figure 4:**
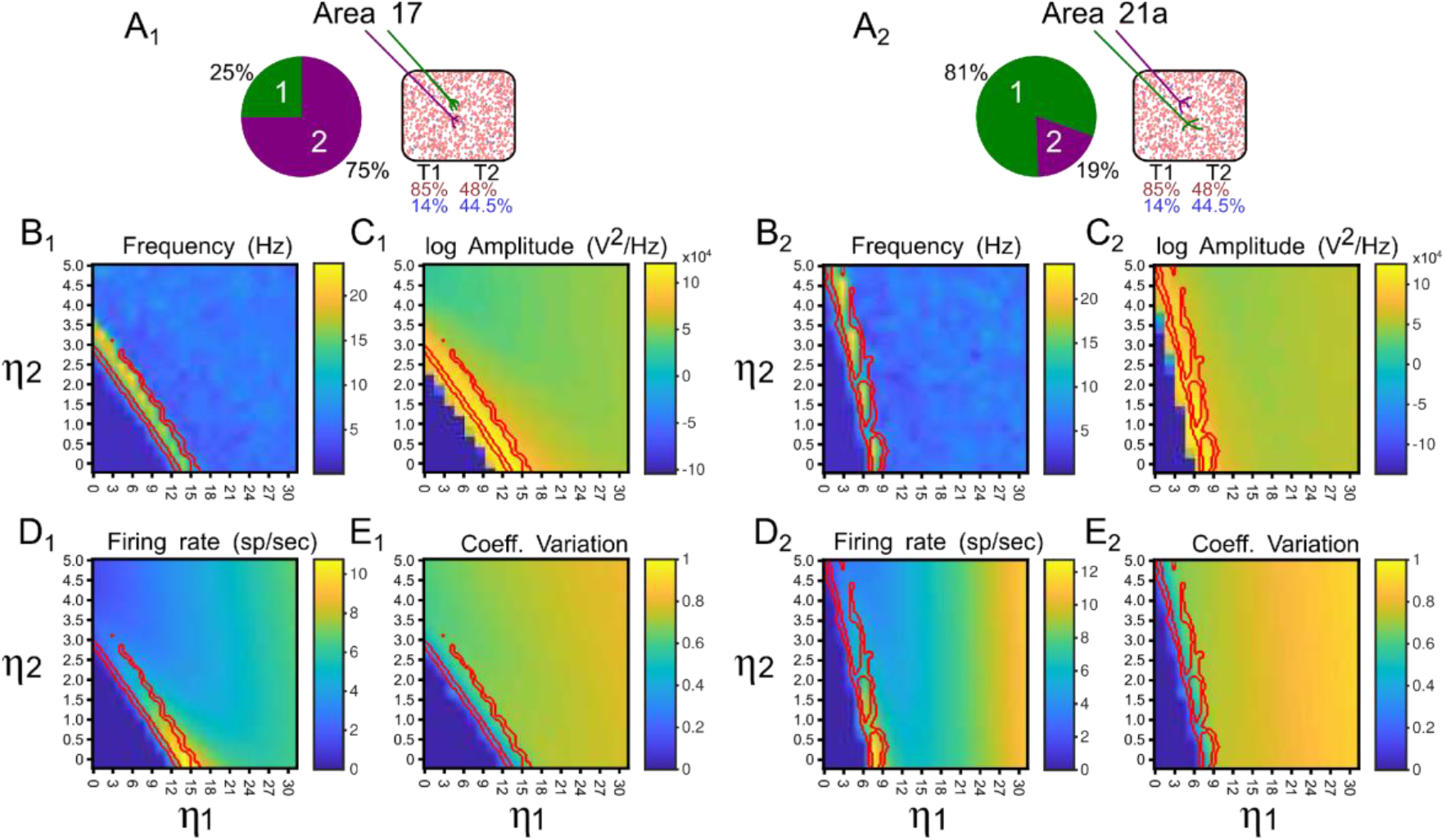
Quantitative outputs for a pulvinar network whose neurons receive projections from areas 17 (1) and 21a (2). A) Distribution and number of contacts to excitatory and inhibitory pulvinar neurons of terminal types 1 and 2 for cortical areas, B) Frequency (Hz), C) Amplitude (v^2^/Hz), D) Firing rate (sp/sec), E) CV.

#### Effect on alpha waves of sequential activation of two cortical areas

Here, alpha-band oscillations generated by the driven area were measured when the other area was activated afterwards. To elicit alpha rhythms that were representative of cortical areas 17 and 21a, we selected strengths of their connections such that *η*_2_ > *η*_1_, and *η*_1_ > *η*_2_, respectively. In this setting, the thalamic network was simulated for 5 sec. After the network reached a stable alpha-band oscillation induced by the driven area (2 sec), the other cortical was “attached” (1 sec). Subsequently, the attached area was disconnected, and a recovery period was allowed (2 secs). The results of such simulations are showed in Figure 5. When driven inputs were from area 17, alpha waves raised quickly (~150 ms) inside the network, having a stable oscillatory profile before the end of the first second. The amplitudes of such oscillations were low (~15 sp/bin), even if pulvinar neurons were synchronized. In the attached period, inputs from area 21a abolished the synchronization, including alpha rhythms. Once 21a was disconnected, the pulvinar network came back rapidly to alpha-band oscillations again. When the driven input was from area 21a, the rise of alpha waves was much slower (~1000 ms), but amplitudes of the oscillation were much larger (~100 sp/bin). Adding projection from area 17 induced an asynchronous stable activity on the thalamic population, which returned quickly to alpha waves when this attached area was disconnected. In summary, the pulvinar network developed alpha rhythms by drivining cortical inputs and, at these parameter values, the arrival of other cortical sources generated a global asynchronous state and a loss of oscillatory alpha-band activity.

**Figure 5:**
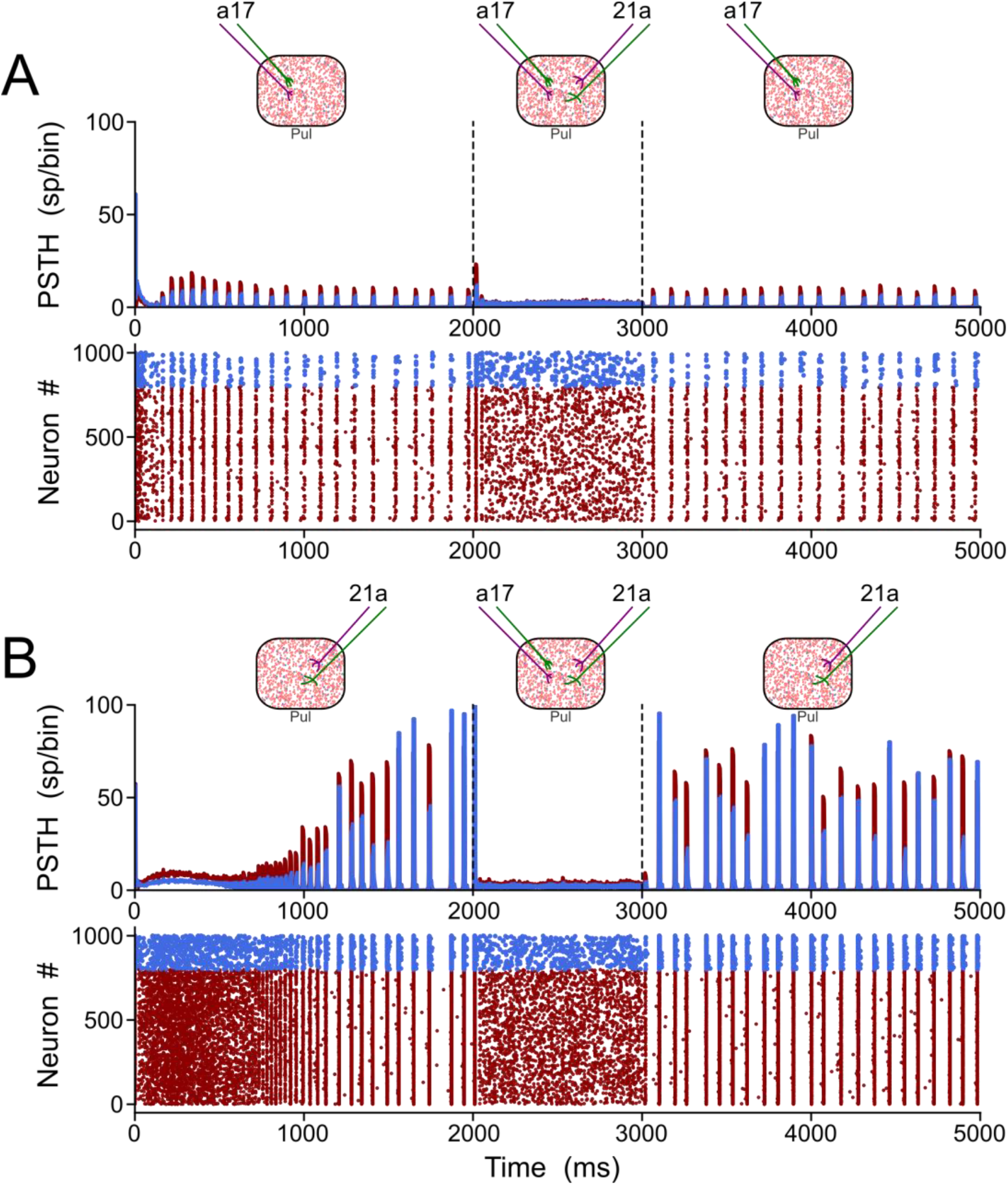
Two solutions to generate alpha waves in the pulvinar. These solutions are found when *η*_1_ > *η*_2_ or *η*_1_ < *η*_2_ are settled for areas 17 and 21a, respectively. Note that the thalamic alpha rhythms are ceased when the other area functionally target the pulvinar (2-3 secs). Solution for areas A) 17 and B) 21a.

Figure 5 showed that terminal types 1 and 2 seem to produce different activation dynamics in the thalamic network, particularly at the beginning of the stimulation. As previous pulvinar responses were only analyzed after 1 second of “recording”, the aim of the following section was to quantify pulvinar dynamics just before the onset of cortical stimulation. For this purpose, *η*_1_ and *η*_2_ were gradually increased for the strength of projections from areas 17 and 21a. The result of such iterations revealed that pulvinar alpha waves were located in similar activation zones shown above (Figure 6). In this regime, pulvinar alpha rhythms appeared and stabilized rapidly when type 2 terminals dominated cortical projections of area 17 (Figure 6A1-A2). Conversely, adding type 1 terminals and decreasing the strength of type 2 axons restricted such oscillations to a short time window (Figure 6A3). In fact, increasing the pathway strength of only type 1 axons into the pulvinar network, generated an asynchronous transition of spike activity which was subsequently transformed into a synchronous oscillation when *η*_1_ was large (Figure 6C1-C3). These qualitative details were further analyzed by measuring averages dispersion (CV), number and the first-time onset of the PSTH peaks of the oscillatory pulvinar synchrony (Figure 6B-D). For area 17, alpha-waves were highly regular (low CV) when *η*_2_ > *η*_1_ (Figure 6B1), with a constant number of peaks and rapid triggering of activity in similar regimes where alpha rhythms were stable over longer simulation times (Figure 6B2-B3). Alpha rhythms for area 21a were also expressed in the same regime (*i.e*., *η*_2_ > *η*_1_). Here, however, when *η*_2_ < *η*_1_, alpha waves were less evident and stable oscillations with regular periodicity in the first second of simulation were undetected. Thus, in the first period of cortical stimulation, type 2 terminals were more likely to establish effectively a stable and fast alpha-band periodicity than type 1 axons regardless of their source area.

**Figure 6:**
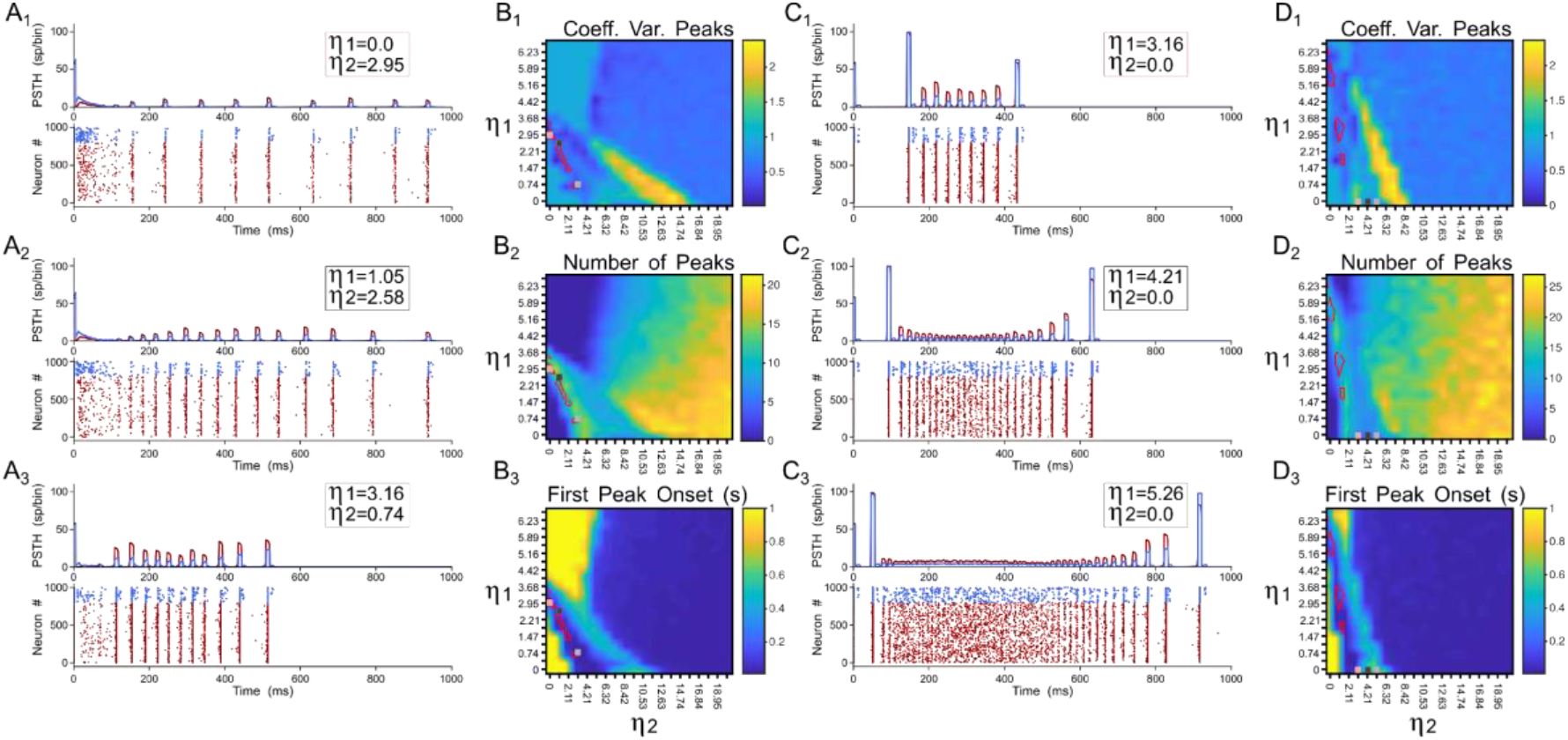
Terminal types 1 and 2 produce different activation dynamics in the thalamic network at the beginning of the stimulation. A) PSTH and raster plots for area 17 solutions. B) Outcomes of such solutions for 1) coefficient of variation of alpha wave peaks, 2) number of peaks per second, and 3) first peak onset. C) PSTH and raster plots for area 21a solutions. D) Outcomes of such solutions for 1) coefficient of variation of alpha wave peaks, 2) number of peaks per second, and 3) first peak onset.

#### Simultaneous cortical activations

The generation of alpha waves in the pulvinar was studied when the two cortico-thalamic projections were combined. The objective here was to evoke alpha waves in the pulvinar with the two cortical areas activated simultaneously. This joint activation may be possible because, for instances, the superposition of oscillatory outputs from areas 17 and 21a (matrices from Figures 3B1 and 3B2) generates spots of alpha-wave responses. To that end, the connectivity weights were iterated to find representative examples of cortico-pulvinar projections that achieved alpha rhythms in the thalamus. Such results are shown in Figure 7, in which the driver activity to pulvinar neurons was originated from the simultaneous convergence of the two cortical sources. In panel 7A, the two cortical areas had weights of type 2 connections weaker than those from type 1, whereas, in panel 7B, terminal types 1 and 2 from area17 were stronger than those from area 21a. The convergence of the two cortical inputs yielded a stable oscillatory alpha-band response after 1 second, which ended when such projections were disconnected (after 2.5 seconds). To demonstrate that the oscillatory activity was evoked by the synergy of the two cortical areas, the thalamic network was initially connected by only one cortical area. After a period (1.5 sec), the input from the other cortical area was restored (Figures 7A2-A3 and B2-B3). Terminal weights used here were the same as before. In the two above cases, the input from one area was insufficient to gate oscillatory responses in the pulvinar (Figure 7A2-A3 and B2-B3). When the weight of connections from area 17 to the pulvinar were more robust than those weights from area 21a (Figure 7B3), thalamic neurons showed oscillatory responses, but only during a short period (<1.0 sec). In this regime of connections, alpha waves in the pulvinar recovered when connections from the disconnected cortical were added from area 17 or area 21a, showing a stable periodicity during the whole period of dual cortical stimulation.

**Figure 7:**
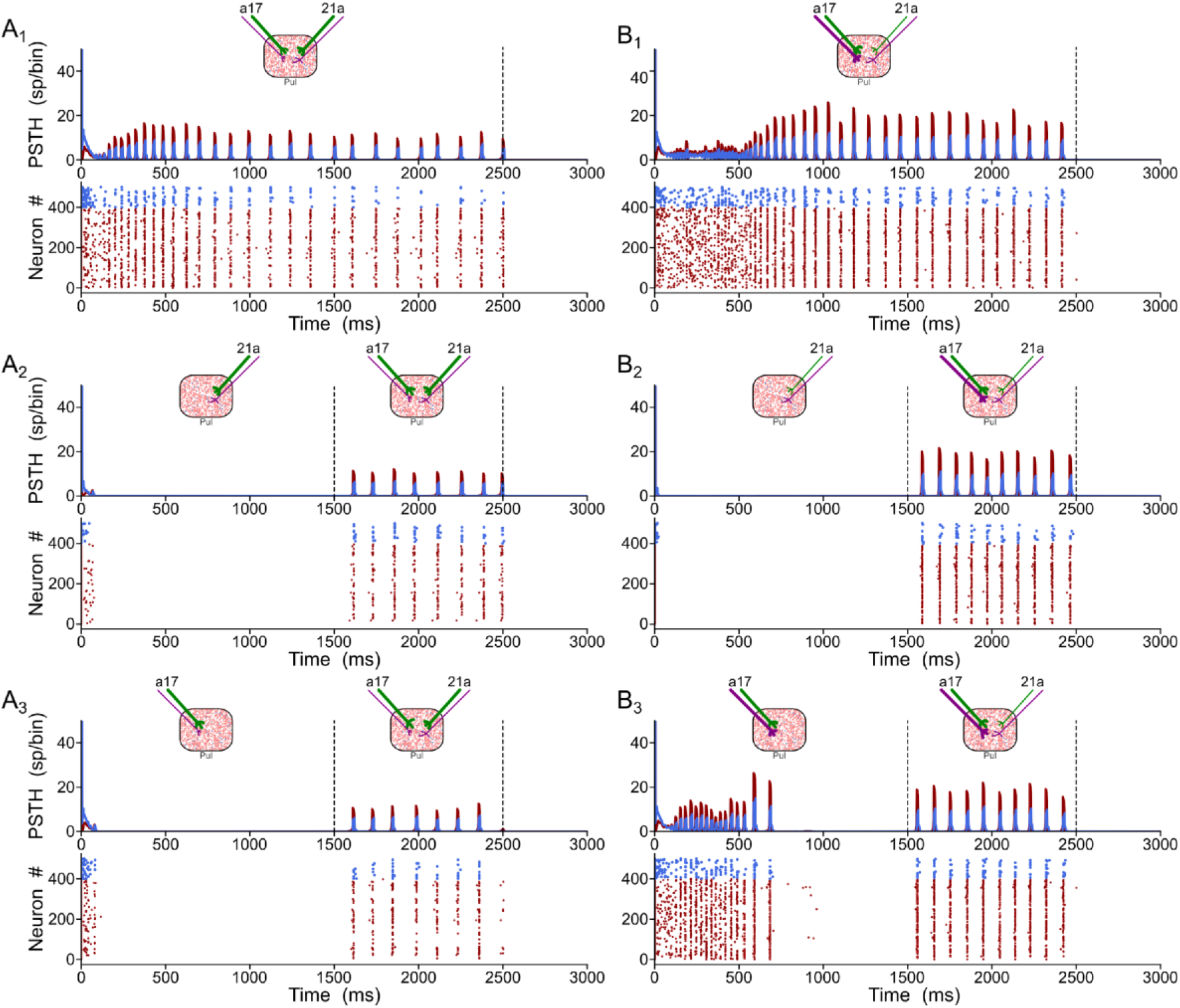
Generation of alpha-waves by the combination of the two cortico-thalamic projections. A) Solution when type 2 terminals (purple axons) are weaker than type 1 (green axons). B) Solution when terminal types 1 and 2 from area 17 are stronger than connections from area 21a.

### 3.4 Oscillatory Responses of the Transthalamic Network

In this section, each cortical was modelled explicitly as a network of *E-I* neurons whose responses were settled in the balanced state. Randomly chosen excitatory neurons formed the two corticothalamic pathways from areas 17 and 21a (*p* = 0.2, homogeneous distribution). The proportion of terminal types 1 and 2 were 2% and 75% and 81% and 29% of *Np* = *K* synapses, for areas 17 and 21a, respectively. While the proportion of excitatory type 1 terminals contacting excitatory and inhibitory pulvinar neurons was *p* = 0.85 and *p* = 0.14, respectively, for type 2 terminals it was *p* = 0.48 and *p* = 0.445, respectively. The two cortical areas were also connected feedforwardly through randomly chosen excitatory connections from area 17 to excitatory and inhibitory neurons of area 21a (*p* = 0.05). Inputs to area 17, from an implicit lateral geniculate thalamic nuclei (LGN), were also drawn from a uniform distribution (*p* = 0.05), and were modelled as spike trains with Poisson statistics. Axon terminals from LGN to area 17, and from area 17 to area 21 did not have any synaptic plasticity. The three interconnected structures mimic the cortico-thalamo-cortical network formed by areas 17 and 21a and the pulvinar.

#### 3.4.1 Effect of Cortical and Pulvinar Dynamics in Oscillatory Responses of the Transthalamic Network

Burst discharges and low background levels of synaptic noise in the pulvinar were first settled to study the cortico-pulvinar pathway of the model. It has been postulated that intrinsic burst neurons in layer 5 (5IB) of the lower cortical areas (*i.e*., areas 17 and 18) participate in the conduction of pulvinar alpha rhythms [O’Reilly et al., 2021]. Furthermore, thalamic bursting has been characterized as an intrinsic property of pulvinar neurons [Ramcharan et al., 2005]. Besides, background noise can alter the synaptic efficiency of connections by increasing subthreshold fluctuations [Silver, 2010] and enhancing the salience of neuronal oscillations to avoid long asynchronous/synchronous state transitions as in Figure 6C1-C3. For that end, intrinsic bursting was induced in cortical and pulvinar neurons by changing mainly high reset values (*V_r_* > *V_T_*), among other parameters of the current regular spike train [Brette and Gerstner, 2005]. Furthermore, stochastic balanced synaptic inputs without short-term plasticity were added equally to *E-I* pulvinar neurons to elicit background noise. Thus, the effect of cortical burst spikes and background noise into pulvinar neurons on thalamic alpha-waves were compared to cases where no such functional biological processes were present.

Figure 8 shows all cases of combinations where area 17 and pulvinar neurons have regular or burst spiking discharges and pulvinar is with or without background synaptic noise. Raster plots for cortical and pulvinar neurons (top and bottom panels, respectively) and PSTH envelopes for excitatory and inhibitory pulvinar neurons (middle panel) are showed here. To obtain such results, cortical area 17 was targeted by an LGN input of 50 sp/sec, and cortical and pulvinar excitatory and inhibitory neurons were recorded for 1 sec. Parameters of the brain structures were fixed between simulations, but cortico-pulvinar weights were fitted to obtain pulvinar alpha rhythms. Cases with background noise in the pulvinar were analyzed first, then when neurons from area 17 had bursting discharges, and finally, when thalamic neurons incorporated such burst-like dynamics. Adding background noise to pulvinar excitatory and inhibitory populations of neurons improved the profile of alpha-band oscillations (Figures 8A - 8B). The average dispersion of the excitatory bumps in the PSTH formed around the synchronous discharge of neurons were 30.05 ± 0.14 ms and 32.54 ± 0.42 ms for simulations with or without background noise in the pulvinar, respectively. When cortical neurons had bursting dynamics (Figures 8C - 8D), bumps of alpha-waves in the pulvinar with and without background, became larger with higher peaks (~10 sp/bin) and smaller dispersion (~29.05 ms), compared to previous cases with only regular spiking discharges. Changing pulvinar regular spiking cells with bursting neurons favored the irregular periodicity of alpha rhythms, but background noise added into excitatory and inhibitory populations stabilized partially such oscillatory lost (Figures 8E - 8F). Taken together, synchronous alpha rhythm responses of pulvinar neurons were affected by cortical and pulvinar neuronal dynamics as well as the background noise added to thalamic neurons.

**Figure 8:**
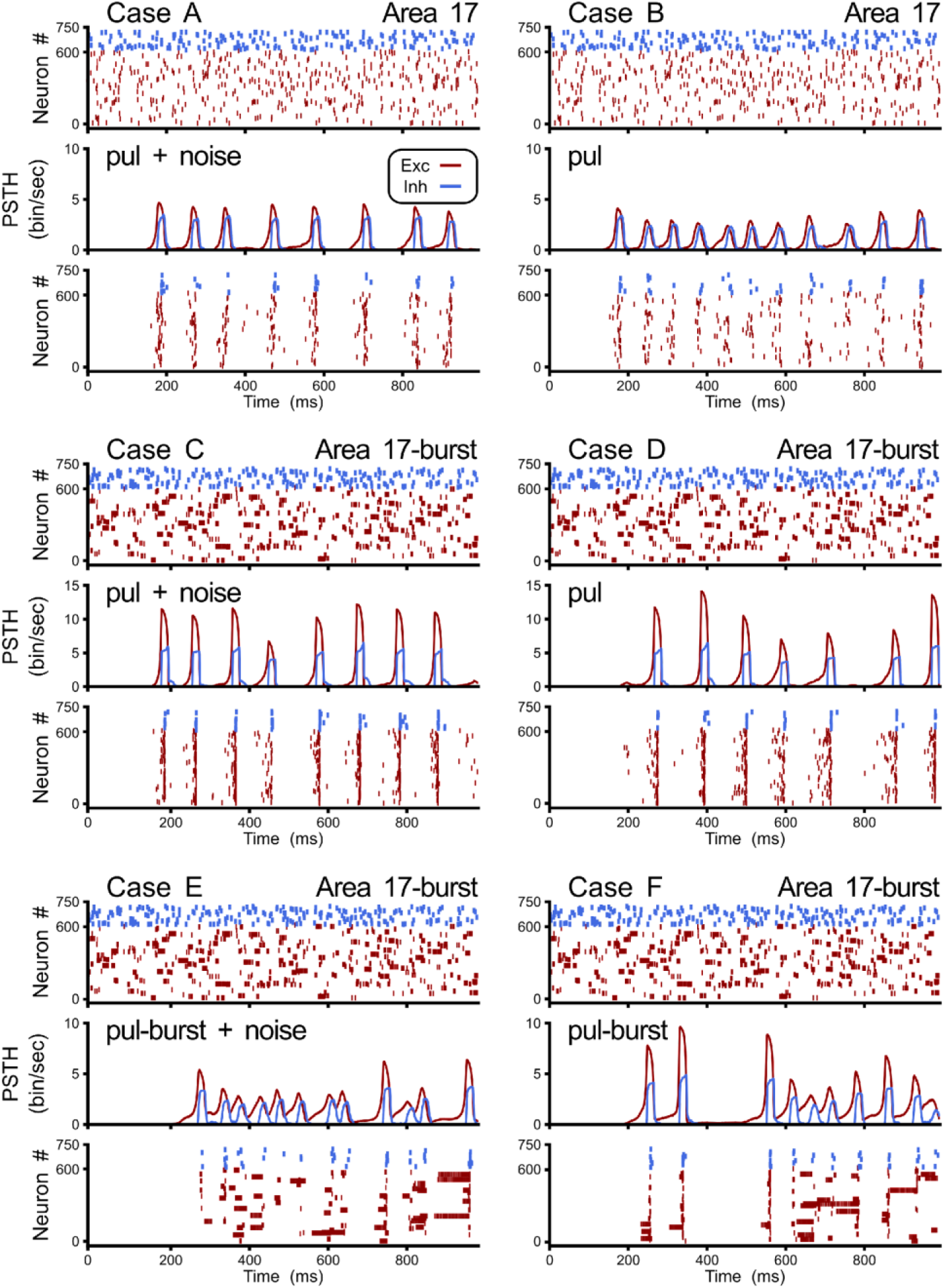
Combinations of cases when area 17 and pulvinar neurons have regular or burst dis-charges, and pulvinar is with or without background synaptic noise. For each case, raster plots for area 17 and pulvinar and PSTH for pulvinar are shown. Case A) Regular spiking for area 17 and pulvinar neurons, background synaptic noise added to pulvinar neurons. Case B) Similar than case A, but without background synaptic noise in pulvinar. Case C) Bursting dynamics for area 17, regular spiking for pulvinar neurons, background synaptic noise added to pulvinar neurons. D) Similar than case C, but pulvinar neurons without background synaptic noise. E) Bursting dynamics for area 17 and pulvinar neurons, background synaptic noise added to pulvinar neurons. F) Similar than case E, but pulvinar neurons without background synaptic noise.

Pulvinar responses to different cortical and pulvinar dynamics were further investigated by measuring oscillatory frequency and amplitude, as well as firing rate and CV when *η*_1_ and *η*_2_ were iterated (Figure 9). Alphawaves were highlighted by depicting contours that had high and low amplitude spectral density in the Fourier transform (Figure 9, red and white lines, respectively). In general, alpha waves were between the transition of quiescent states and stable asynchronous activity. While strong alpha-band oscillations were limited to be in such transition (red lines), low energy frequencies were revealed to be more ubiquitous (white lines). Noise decreases the weights used to evoke such strong alpha-frequency oscillatory responses, when comparing iterations with and without background noise (Figure 9, A vs B and C vs D). Background noise also restricted the feasible region for finding the alpha waves. On the contrary, incorporating cortical bursting dynamics in the network expanded such regions, particularly for weak alpha-band oscillations (Figure 9B-E). Strong alpha rhythms were eliminated when burst responses were included into pulvinar neuron dynamics. As figures 9D-E show, weak alpha-waves were localized at the transition from silent to activated states, when background synaptic noise was present and cortical and pulvinar neurons had bursting dynamics. Taken together, incorporating bursting cortical and pulvinar dynamics and adding background synaptic noise into pulvinar neurons enhanced oscillatory alpha-band states of the cortico-pulvinar network, but the localization of such low-frequency transition regimes was almost unchanged.

**Figure 9:**
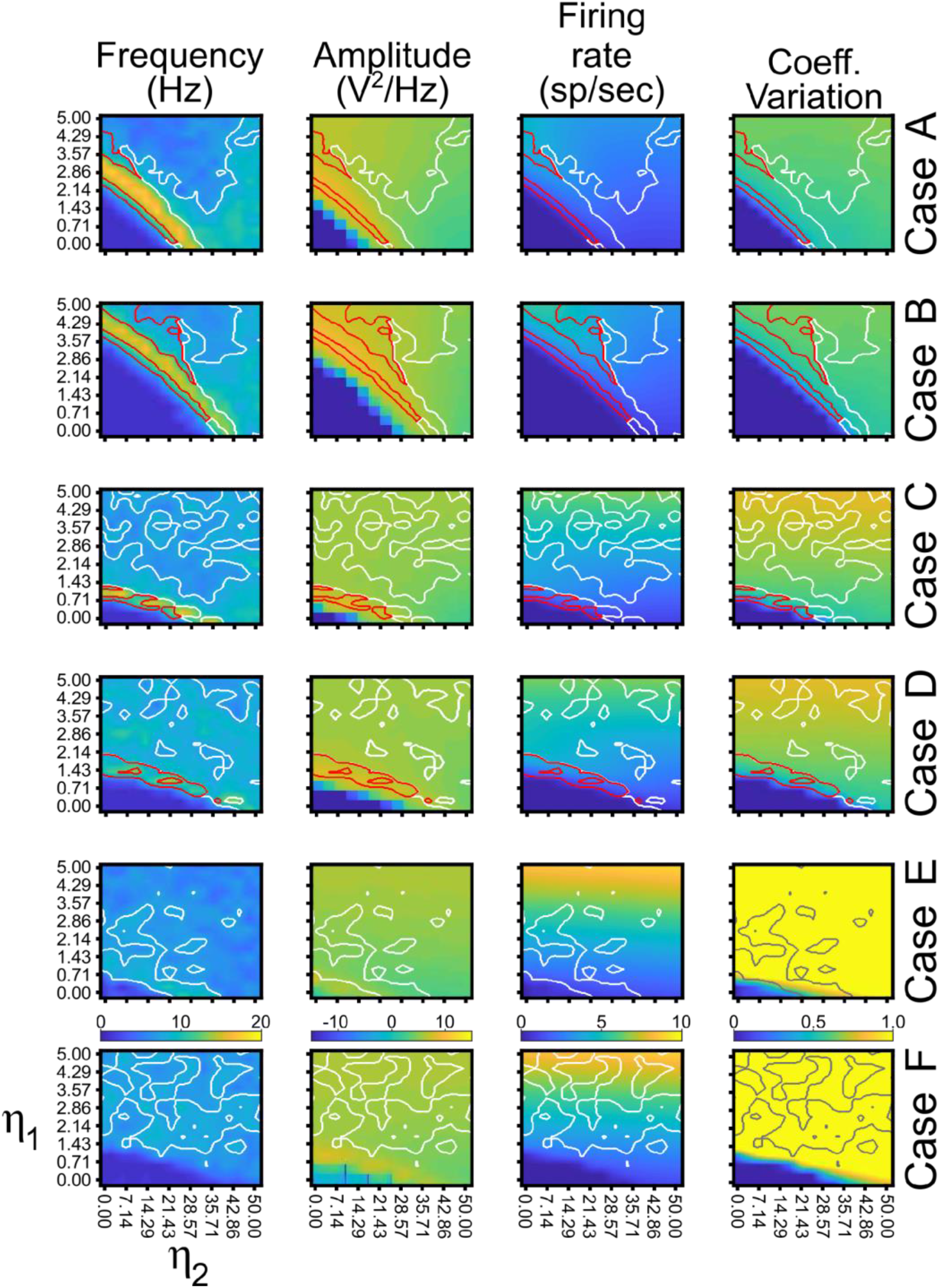
Quantitative outputs for cases in Figure 8. Frequency (Hz), amplitude (v^2^/Hz), firing rate (sp/sec) and CV for different cases as *η*_1_ and *η*_2_ increase gradually. Red line, alpha-waves with higher amplitude; white line, alpha-waves with lower amplitude.

#### 3.4.2 Effect of Tranthalamic Pathway in Higher-Order Cortical Responses

Pulvinar effective connectivity to the visual cortex was tested by forming the transthalamic pathway. In this scenario, cortical area 21a received two feedforward projections: one coming from area 17 and another from the pulvinar (Figure 1C3). In turn, area 17 received an input from the LGN that consisted of spike trains with Poisson statistics. Connections from the LGN to area 21a were not implemented. The pulvinar was divided in two functional areas based on the arrangement of cortico-thalamic projections: striate- and extrastriate-recipient zones. Each recipient zone received percentage of connections and excitatory and inhibitory proportion of contacts from terminal types 1 and 2 of cortical areas 17 and 21a as previous simulations. However, connections between *E-I* pulvinar cells were strong, similar to the organization of the two cortical areas. For simplicity, only one pulvino-cortical projection was considered, which was for the connection from the pulvinar to area 21a. For this area, each excitatory and inhibitory populations received in average *K* random contacts from excitatory pulvinar neurons. Weights of this pulvino-cortical projection, *G*^21*a*←*pul*_*E0*_^ and *G*^21*a*←*pul*_*I0*_^, were multiplied by the same factor, *W_CP^21a^_*, which was fixed for all simulations. Dynamics of the pulvinar neurons used here were those established in case A (Figure 9A), when the neurons had regular spiking and low background synaptic noise. Thus, the neural network used for the next iterations consisted of a cortico-cortical feedforward pathway from area 17 to area 21a, and a transthalamic pathway from area 17 to the pulvinar, and from pulvinar to area 21a, also defined by cortico-thalamic projections from areas 17 and 21a to striate- and extrastriate-recipient zones in the pulvinar, respectively.

##### Propagation of alpha rhythms from pulvinar to area 21a

To achieve alpha waves in the pulvinar, cortico-thalamic projections from area 17 to the pulvinar were fixed to a constant value, and cortico-thalamic weights from area 21a to the pulvinar were gradually varied. Figure 10 shows an overview of the network performance. For these simulations only, *η*^21*a*_*1*_^ = *η*^21*a*_*2*_^. During the first second, the cortico-cortical pathway was attached, whereas the cortico-thalamo-cortical pathway was unconnected. By consequence, predominant low-frequency oscillations were absent in area 21a. Then, cortico-thalamic from area 17 and pulvino-cortical projections to area 21a were added. In this configuration, alpha-band oscillations were generated in cortical area 21a by the transthalamic pathway (1-2 secs). Note that during this period, alpha-waves were only evoked in the pulvinar striate-recipient zone. In the next temporal sequence (2-6 secs), cortico-thalamic connections from area 21a were added into the pulvinar extrastriate-recipient zone, and these projections were amplified gradually throughout this period. Alpha waves in area 21a and the pulvinar persisted when cortical projections had a low magnitude (2-3 secs). In the contrary, alpha-band oscillations were lost when *η*^21*a_1_*^ and *η*^21*a_2_*^ enhanced (3-6 secs). The pulvinar and area 21a showed an asynchronous profile of spike activity when cortical top-down connections to the pulvinar were sufficiently high. Note that pulvinar responses were produced only in the extrastriate-recipient zone, while responses in the striate-recipient zone were silenced. Finally, alpha-waves in the pulvinar and area 21a were re-established when cortico-pulvinar connections from area 21a were disconnected (6-7 secs). Taken together, the transthalamic pathway enabled functional connectivity in alpha frequency band when cortico-pulvinar connections from lower cortical areas were allowed and an asynchronous response in higher cortical areas when the cortico-pulvinar loop between area 21a and pulvinar was closed.

**Figure 10:**
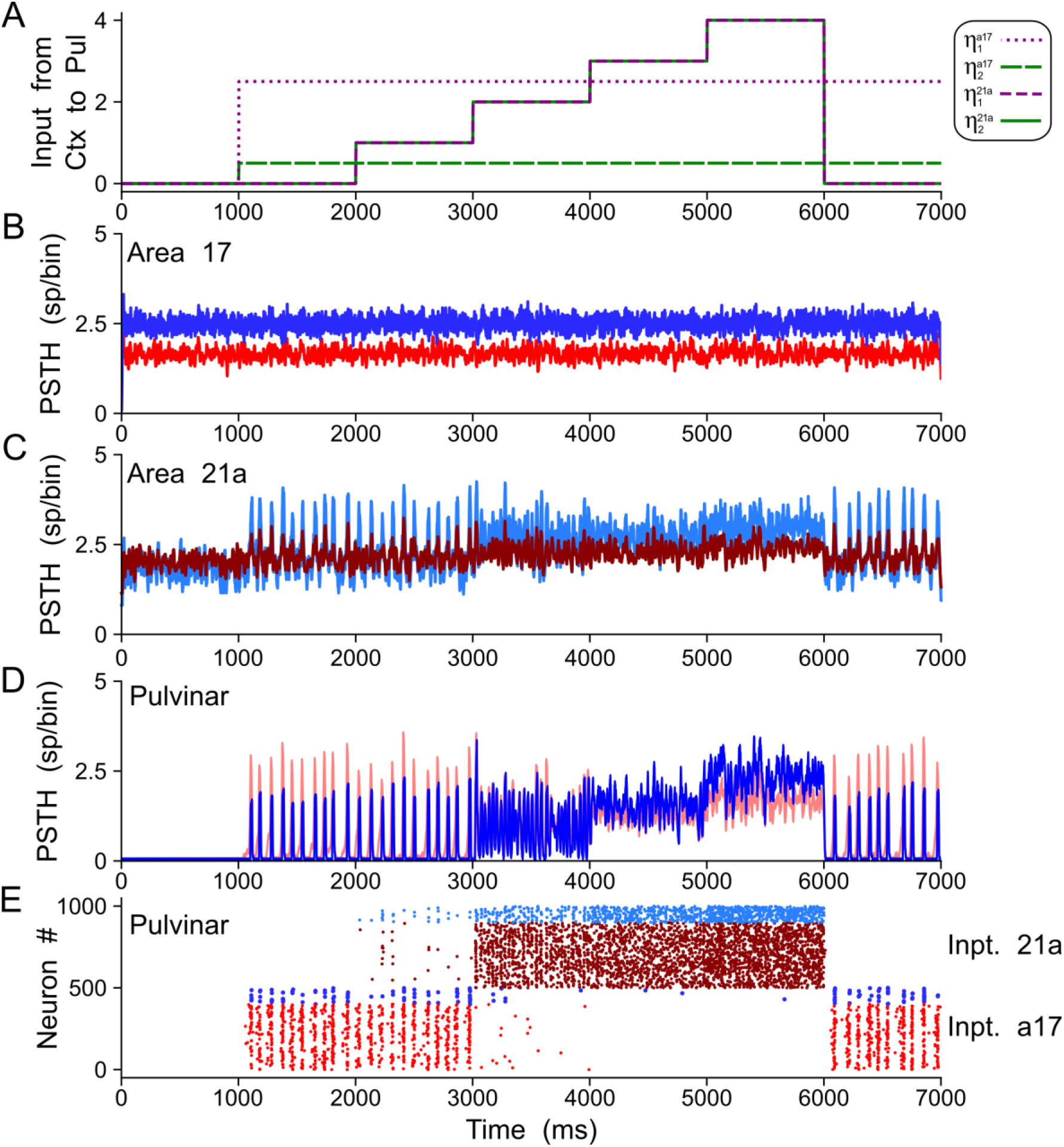
Example of oscillatory alpha-band activity transmission by transthalamic network for a network as Figure 1A3. A) Weights of connections, *η*^*a*17,21*a_1_*^ and *η*^*a*17,21*a_2_*^ for areas 17 and 21a. PSTH for areas 17 (B), 21a (C) and pulvinar (D). E) Raster plots for neurons in the pulvinar. Note that half of the pulvinar neurons are target by cortico-thalamic projections from area 17 and the other from area 21a.

##### Closing the cortico-pulvinar cortical loop

The effect of closing the loop between cortical area 21a and the pulvinar on the formation of alpha waves was further analyzed by varying the weights of terminal types 1 and 2 of these cortico-thalamic projections. For that end, a network incorporating cortico-cortical and transthalamic pathways (area 17 to pulvinar, pulvinar to area 21a) was created as above described. In addition, the network had a top-down projection from area 21a to the pulvinar, which was organized in three different time periods. In the first condition, the projection from area 21a and the pulvinar was disconnected (open loop) for 1 sec (Figure 11A). The next second, the projection from area 21a to the pulvinar was formed (close loop), and connections from area 17 to the pulvinar were still activated. In the last second, only the close loop was functional, in which cortical top-down activity to the pulvinar was evoked indirectly by cortico-cortical arriving inputs to area 21a (Figure 11B).

**Figure 11:**
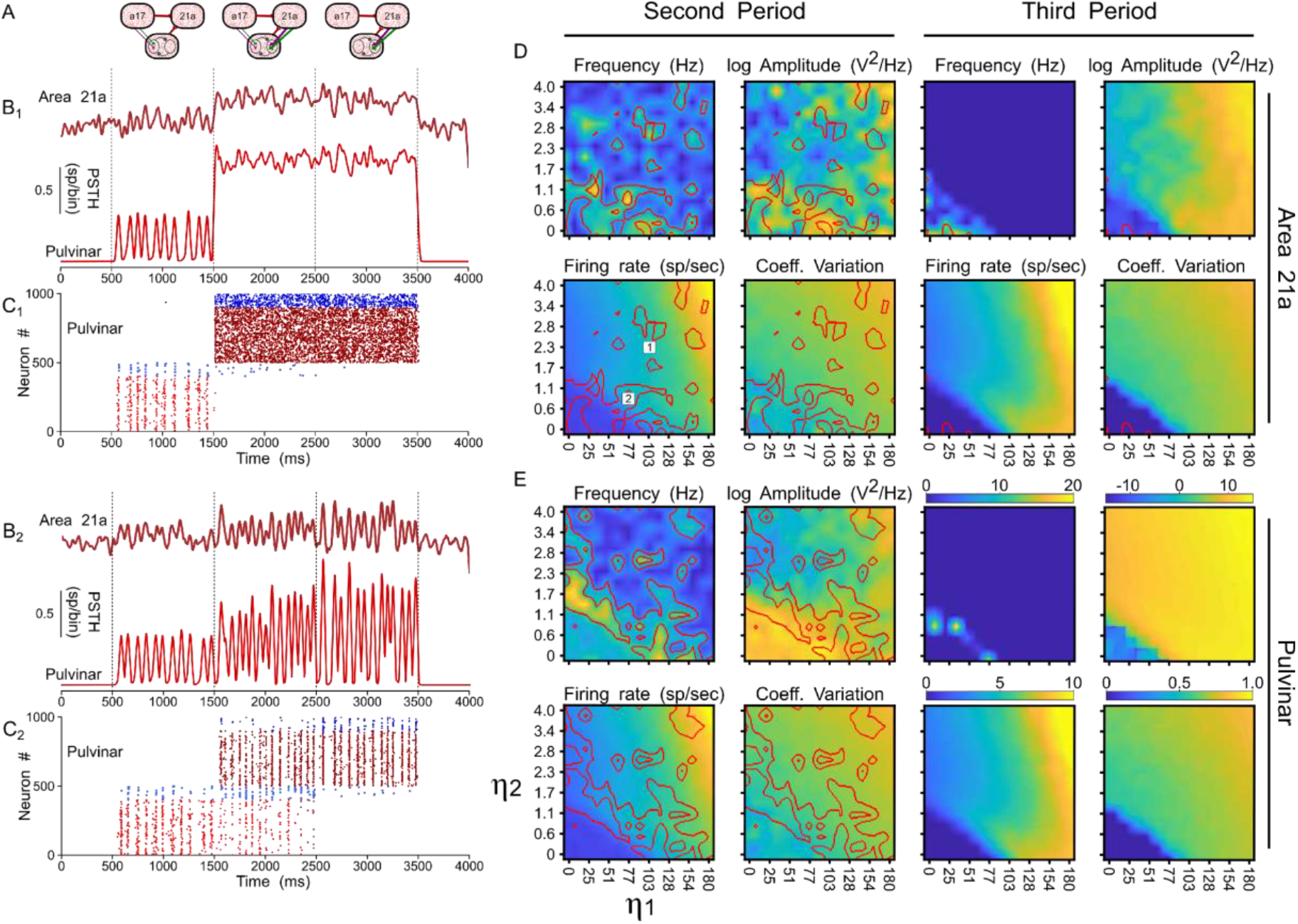
Effects on alpha waves when the loop between area 21a and the pulvinar is closed in the cortico-thalamo-cortical model. A) Scheme showing the temporal configuration of the three periods for the connections of the model. Panels B) and C) are two solutions when topdown loop to the pulvinar generate asynchronous (1) or oscillatory (2) responses. B) PSTH for area 21a and pulvinar. C) Raster plots for pulvinar neurons with striate- and extrastriate-recipient zones. Second and third periods’ quantitative outputs (Frequencies, amplitudes, firing rates and CVs) for area 21a (D) and (C) pulvinar networks when *η*^21*a_1_*^ and *η*^21*a_2_*^ are increased gradually. Note the position of solution 1 and 2 in the panel D.

With this network, the efficacy of alpha waves expressed in area 21a by pulvinar projections was quantified. For the first second, alpha waves in the pulvinar and area 21a were evoked 99% and 64.44% of the time, respectively. The remaining ~35% of oscillations evoked in area 21a were lower than 7.5 Hz. In the next condition, *η*^21*a_1_*^ and *η*^21*a_2_*^ were iterated and the frequencies, amplitudes, firing rates and CV of area 21a and pulvinar outputs were characterized. Figure 11C shows that, in the second period (2-3 secs), closing the loop between area 21a and the pulvinar decreased alpha-band oscillations in the two networks. In fact, the pulvinar achieved only 36% of alpha-waves, causing a loss of alpha-wave transmission in area 21a (~26%). Alpha-band oscillations in the pulvinar were only expressed at low magnitudes of *η*^21*a_1_*^ and *η*^21*a_2_*^, since the striate-recipient zone continued to yield such rhythms. When the strength of the cortico-thalamo projections, from area 21a to the pulvinar extrastriate-recipient zone was increased slightly more, the striate-recipient zone was perturbated, and so was the production of alpha-waves, decreasing the propagation of such frequency into area 21a during this second period. Note that a variety of higher frequency oscillations in area 21a and the pulvinar arose at such intermediate neuronal states (Figure 11B1). As the forces of the cortico-thalamic connections were higher, only extrastriate-recipient neuronal responses in the pulvinar were engaged, causing an asynchronous spiking state in both the pulvinar and area 21a (Figure 11B2). Similar responses were observed in the last period of stimulation (3-4 secs). Here, the close loop was still coupled, but connections from area 17 to the pulvinar were disconnected. Under this configuration, only 1.7% of alpha waves were generated in area 21a. In average, the pulvinar did not show any alpha waves during this period. The remaining spiking responses of area 21a and the pulvinar had almost all asynchronous features or higher-frequency oscillations (Table 2). Altogether, similar to previous analysis, the transthalamic pathway allowed a propagation of alpha waves from the pulvinar to area 21a, which was occluded by the arriving of the top-down projection from area 21a to the pulvinar. Thus, the pulvinar showed a bi-stable spiking response, oscillatory or irregular spiking, whose responses were gated by the different activations of projections from areas 17 and 21a to the pulvinar.

**Table 2:** Percentage of low-frequency oscillations for pulvinar and area 21a, during the 3 recording periods of Figure 11.

|  | Alpha (%) | >Alpha (%) | <Alpha (%) | Period |
| --- | --- | --- | --- | --- |
| Pulvinar | 99.03 | 0.78 | 0.14 | 1 |
|  | 35.97 | 10.95 | 52.87 | 2 |
|  | 0.00 | 0.00 | 100.00 | 3 |
| Area 21a | 64.44 | 0.73 | 34.54 | 1 |
|  | 26.06 | 1.89 | 71.70 | 2 |
|  | 1.69 | 0.03 | 98.29 | 3 |

## 4 Discussion

Previous experimental studies have shown that neuronal populations of the pulvinar nucleus possess multiple dynamic activity profiles [Wrobel et al., 2007, Saalmann and Kastner, 2011, Yu et al., 2018, Le et al., 2019]. In this work, using a theoretical approach, we propose that such population dynamics in the pulvinar may arise from cortico-thalamic connections originating from different hierarchical cortical areas, whose axonal terminals have distinct anatomical and physiological profiles. Two types of neuronal discharges were evoked in the pulvinar, synchronous and asynchronous responses, when projections from areas 17 and 21a to the pulvinar were organized as the connection empirically described in cats [Huppe-Gourgues et al., 2006, 2019, Abbas-Farishta et al., 2020]. We found that, at a specific connection weights regime, cortico-thalamic connections reproduce the oscillatory alpha-band activity of the pulvinar. This solution from the model was found when each cortical area independently contacted the pulvinar and when they were combined. When a single area evoked oscillatory alpha-band activity in the pulvinar, the activation of the distant area ceased such oscillations. The combinations of the two projections changed the oscillatory dynamics to asynchronous spiking states. These bi-stable states may have implications for the propagation of visual responses along the cortical hierarchy. The results found in our models are a direct consequence of the restrictions we put on the parameters, suggesting that the pulvinar may have several dynamic response states to interact with the visual cortex.

The most important prediction of this work is that the pulvinar has at least two functional response states, regular oscillatory or stable asynchronous. Other models explaining such oscillatory variations have included alpha waves in the pulvinar implicitly [Quax et al., 2017] or only as part of the cortico-cortical circuitry [Jaramillo et al., 2019]. Our model suggests that part of such oscillatory activity comes from cortico-thalamic projections distributed along the hierarchy of the visual system. Although alpha rhythms in the pulvinar may reflect the functional cortical feedback connectivity [Halgren et al., 2019], our model predicts that such oscillations may be abolished by cortico-thalamic projections from lower cortical areas (areas 17 and 18). As shown here, given that such connections have a high proportion of type 2 connections, their role would be to desynchronize the pulvinar when activated in a synchronous manner. The implications of such predictions are explained in more detail below.

### 4.1.1 Mechanism of generating oscillations

To obtain oscillatory alpha-band activity, in our model, we settled connections to be unbalanced, so cortico-thalamic weights were just lower than √K synapses providing low-frequency waves. Since oscillatory solutions were in the frontier of a balanced state (Figure 5), the addition of an extra input generated an asynchronous irregular spiking activity. As shown in Figure 6, type 1 terminals generated stable oscillations in late periods, whereas for type 2, the regular periodicity was reached early. The addition of a synaptic noise background (Figure 8A) allowed decreasing the delay to reach this oscillatory stability [Silver, 2010]. The addition of burst-like dynamics favored the appearance of more realistic oscillations than one would expect in an experimental work (irregular oscillations and lower power in the amplitude of the signal) [Ramcharan et al., 2005, O’Reilly et al., 2021]. However, when the cortico-pulvino-cortical model was implemented, pulvinar neurons with regular spiking responses with a low background synaptic noise were selected (case A, Figures 10 and 11). Such properties were chosen to reinforce the regular oscillatory activity of the pulvinar on the neural responses of the visual cortex. Although the burst-like responses may refine the neuronal responses of the pulvinar to a more realistic scenario, our model predicts that such stable oscillatory responses are due to the physiological properties of terminals types 1 and 2 and how these are combined to obtain a nearly balanced state.

### 4.1.2 Implications in the Transmission of Oscillatory Cortical Responses throughout the Transthalamic Pathway

Visual processing in cortical areas follows mostly a hierarchical stream of communication, in which the visual information travels from low to high levels (feedforward pathway), as well as from high to low levels (feedback pathway) [Felleman and Van Essen, 1991, Bullier, 2003]. Oscillatory-band activity has been associated with these anatomical pathways, in which gamma-band oscillations are found in the feedforward route, whereas slow oscillations, alpha/beta waves, are observed in the feedback direction [van Kerkoerle et al., 2014, Bastos et al., 2015]. In our simulations, nonetheless, we show that the alpha-type oscillatory activity evoked in area 21a, we are able to reverse that transmission in feedback action if *η*^21*a_1_*^> *η*^21*a_2_*^, *so* reversing the ratio of type 1 and 2 terminals, and introducing a pulvino-cortical projection such as *W_CP_^a17^<0*. In this new scenario, cortico-thalamic projections from area 21a would drive alpha-band oscillatory responses in the pulvinar, whereas CT connections from area 17 to the pulvinar would shift such responses into asynchronous-type activity (Figure 12).

**Figure 12.**
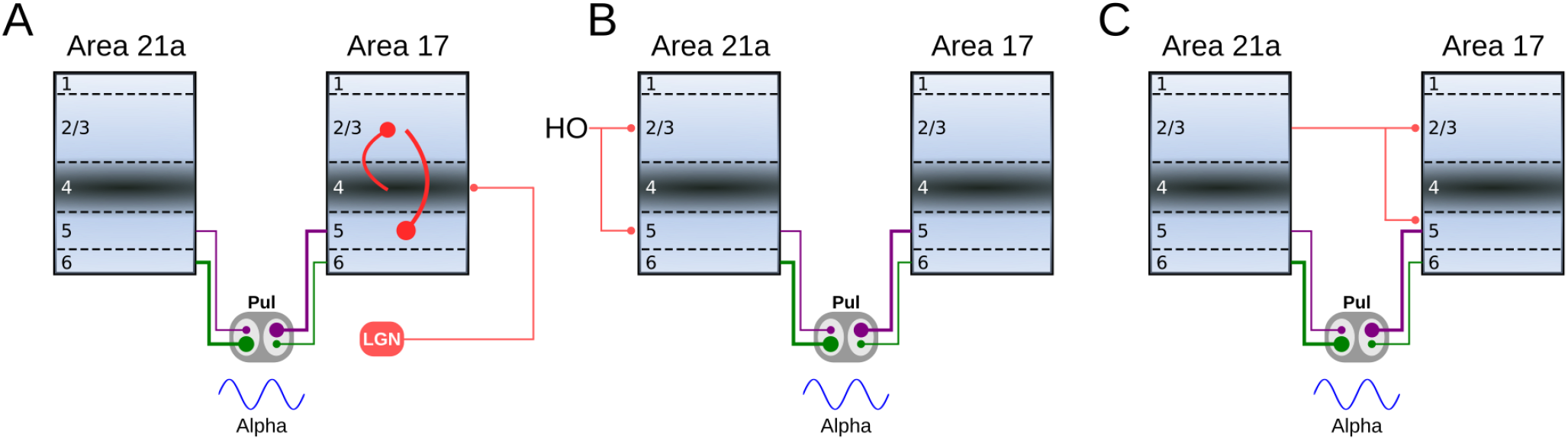
Schematic representation of the oscillatory responses predicted by our models. A) The pulvinar evokes an alpha-band oscillatory response by the feedforward activity of area 17. The response of the infragranular layers of area 17 is evoked by the intralaminar connectivity that initially reaches layer 4. B) Oscillatory alpha-band responses in the pulvinar are evoked by feedback activity from higher-order (HO) cortical areas to the infragranular layers of area 21a. C) Indirect oscillatory alpha-band activation of the pulvinar by the feedback from area 21a to the infragranular layers of area 17.

Reversing the direction of transmission along the cortical hierarchy, and changing the activity from a synchronized to an unsynchronized state, depends on two properties of the simulated network. As described above, terminal types 1 and 2 can evoke alpha-band oscillations independently. Interestingly, as the terminals are set with the ratios of areas 17 and 21a, solutions are found when the predominant terminal had a higher weight of connection than the other type. That is, to engage alpha waves in the pulvinar by cortico-thalamic projections from area 17, the type 2 terminal had to be stronger than the type 1, and vice-versa for area 21a, which follows the ratios found in the empirical anatomical data. The second property is derived of the two compartments organized in the pulvinar as striate- and extrastriate-recipient zones. Our model predicts that activity between these zones would compete with each other for activation in the pulvinar, similar to a “Winner-Take all” (WTA) process. Since these zones are activated independently by different cortical areas along the visual hierarchy, asynchronous or synchronous activity within the pulvinar will also be triggered independently. Such hypothetical compartmentalization of responses predicted by the model allows the pulvinar to revert its oscillatory alpha-band activity to an asynchronous irregular spiking or vice versa. Thus, the pulvinar filters one type of neural response over the other, depending on which cortical area projects more strongly to it.

The pulvinar, across different species, is a heterogeneous structure that is composed of multiple subdivisions and, unlike the LGN, such separations have little organizational arrangements like neuronal lamellae [Baldwin and Bourne, 2017]. These anatomical features mean that the functionality of the pulvinar throughout mammalian evolution remains a mystery [Casanova 2003]. Our model, with at least two-separated cortical-recipient zones, may clarify functional aspects that the pulvinar has. As our model predicts, different low-band frequencies are built in the pulvinar as the force of cortico-thalamic connections increases gradually, mainly when type 1 connections are used (Figures 4 and 11). If several cortical areas are represented independently as separate connections domains in the pulvinar [Shipp, 2003], these pulvinar domains could generate different oscillations regulating visual cortical activity and coordinating transthalamic messages in an oscillatory ascending manner to the visual cortex. Thus, like a WTA computation, the pulvinar would serve as a channel selector of different band frequencies that would adjust cortical dynamics to be able to transmit oscillatory low-frequency activity from one group of neurons to another, possibly in a feedforward or feedback manner throughout the visual hierarchy [Quax et al., 2017, Jaramillo et al., 2019, Cortes et al., 2020]. In other words, the pulvinar could select and separate signals from and to the visual cortex by the above described WTA mechanism. According to this view, the pulvinar would use two anatomical properties to allow visual processing in the cortex: a gradient of terminal types 1 and 2 throughout the cortical hierarchy [Abbas-Farishta et al., 2020], and the spatial processing by the compartmentalization of cortical-recipient zones suggested by our model. Therefore, the pulvinar would need a differential increase in type 1 terminal weights along higher order visual cortical areas and cortico-recipient zones that are partially isolated to carry out this selection mechanism [Huppe-Gourgues et al., 2006, 2019, Abbas Farishta et al., 2020].

The addition of more inputs into the pulvinar would explain the appearance of other low-frequency oscillations observed in experimental data. As our simulations predict, the increase of strengths of the cortico-thalamic projection from area 21a (Figures 3 and 11) causes a shift to other oscillatory frequencies by the competition between rival cortical-recipient zones in the pulvinar One interpretation of these results may be the recruitment of other higher cortical areas that synergistically activate the pulvinar. As more and more areas project to the pulvinar, this top-down activation could generate different band frequencies, which is seen for theta-, beta- and gamma-band oscillations in awake animals [Wrobel et al., 2007, Saalmann and Kastner, 2011, Yu et al., 2018]. On the other hand, the oscillatory response provoked by the joint action of the two areas (Figure 7) could also be interpreted as a convergent action of connections coming from the visual cortex and other brain structures. As our model shows, we found a solution to evoke alpha rhythms when areas 17 and 21a had similar weights of connections. Since the connectivity weights of the cortical areas across the visual hierarchy are unlikely to be similar (because of their different physiological characteristics), one prediction of the model is that one of those connections is not from the visual cortex. Instead, it would arise from subcortical projections, for example, from the superior colliculus. This joint cortical and subcortical action could engage pulvinar responses in different oscillatory modes [Le et al., 2019], when, for example, eye movements are required [Berman and Wurtz, 2011]. Thus, the recruitment of other cortical areas and the combination with other subcortical structures could explain the different oscillatory ranges found in the pulvinar.

Another prediction of our model is that the inhibition of one compartment in the pulvinar can increase the activity in another cortico-recipient zone. For example, in Figure 11, stopping the extrastriate-recipient zone’s activity produces excitation of the striate-recipient zone, which causes an increase of alpha-band oscillations in area 21a by the transthalamic pathway. This excitatory oscillatory effect on cortical populations of neurons when local inhibitions are made in restricted areas of the pulvinar has been already quantified experimentally [Cortes et al., 2020]. Alternatively, this antagonistic effect could also explain the effect of GABA inactivation in the pulvinar, in which the firing rate of cortical neurons achieves a reduction and an enhancement in areas 17 and 21a of cats, respectively [de Souza et al., 2020]. Likewise, the enhancement increase of stimulus-driven responses in V2 cells of monkeys when inactivating the pulvinar [Soares et al., 2004], or even the suppressor effect to the surrounding regions of the receptive fields of V1 neurons when the pulvinar is excited [Purushothaman et al., 2012]. The suggested compartmentalization of the pulvinar would explain why disruption of the activity of isolated domains could create excitation of other nearby domains within the pulvinar.

### 4.1.3 Implications of Pulvinar Bistable States

Perception of low contrast stimuli reveals that the brain shows two temporal states of visual awareness processing [van Vugt et al., 2018]. The brain processes external stimuli in a modular and parallel bottom-up hierarchical fashion if the stimulus is strong enough and exceeds an internal threshold of visual perception. This first state is essentially feedforward. Such signals are retrieved and selected in a second state through attentional requirements located in hierarchically elevated cortical areas. Here, the flow pathway is in a feedback manner, favoring convergence to activate necessarily and sufficiently all the network nodes [Dehaene et al., 2011].

Although both cortical states have been explained based on the long-range excitatory cortical connection, we postulate that the pulvinar is involved in changing from one state to another. Such control would be possible because of the bi-state that the pulvinar has. Theoretically, it has been shown that the pulvinar and its transthalamic pathway are necessary to pass neural responses in a graded manner through a chain of sequentially connected areas [Cortes and van Vreeswijk, 2012, 2015, de Souza et al., 2020]. Such a prediction of the functionality of the transthalamic pathway has been partially corroborated by experiments in the visual system of cats [de Souza et al., 2020]. On the other hand, the differential oscillatory response of the neuronal signals between the cortices in certain phases of pulvinar oscillatory activity [Saalmann and Kastner, 2011, Fiebelkorn et al., 2019], as well as when the pulvinar is inactivated [Cortes et al., 2020], suggest that it plays a role in the effective cortical connectivity. Our current results, showing bi-stable pulvinar states, suggest that cortical feedback transmission is associated with pulvinar oscillatory activity and the feedforward pathway, with the asynchronous irregular spike response of the pulvinar. This prediction of feedforward oscillatory activation should be verified in future works, since pulvinar could be indirectly activated by cortical feedback. That is, from layers 2/3 of a higher cortical area to layer 5 of a lower cortical area, and from here to the pulvinar. Such a feedback pathway would induce the type of antagonistic dynamics predicted here (Figure 12).

## Author Contributions

Conceptualization, N.C.and R.A.F; methodology, N.C.; software, N.C.; validation, N.C. and C.C.; formal analysis, N.C.; investigation, N.C.; resources, C.C.; data curation, N.C.; writing—original draft preparation, N.C.; writing—review and editing, N.C., R.A.F., H.L., and C.C.; visualization, N.C. and C.C.; supervision, C.C.; project administration, C.C.; funding acquisition, C.C. All authors have read and agreed to the published version of the manuscript.

## Acknowledgments

This work was supported by a Canadian Institute for Health Research (CIHR) grant to C.C. (PJT-148959). We thank Lamyae Ikan for her useful discussion.

